# eIF3 associates with 80S ribosomes to promote translation elongation, mitochondrial homeostasis, and muscle health

**DOI:** 10.1101/651240

**Authors:** Yingying Lin, Fajin Li, Linlu Huang, Haoran Duan, Jianhuo Fang, Li Sun, Xudong Xing, Guiyou Tian, Yabin Cheng, Xuerui Yang, Dieter A. Wolf

## Abstract

eIF3 is a multi-subunit complex with numerous functions in canonical translation initiation. eIF3 was also found to interact with 40S and 60S ribosomal proteins and translation elongation factors, but a direct involvement in translation elongation has never been demonstrated. We found that eIF3 deficiency reduced early ribosomal elongation speed between codons 25 and 75 on a set of ∼2,700 mRNAs encoding proteins associated with mitochondrial and membrane functions, resulting in defective synthesis of their encoded proteins. To promote elongation, eIF3 interacts with 80S ribosomes translating the first ∼60 codons and serves to recruit protein quality control factors, functions required for normal mitochondrial physiology. Accordingly, eIF3e^+/-^ mice accumulate defective mitochondria in skeletal muscle and show a progressive decline in muscle strength. Hence, eIF3 interacts with 80S ribosomes to enhance, at the level of early elongation, the synthesis of proteins with membrane-associated functions, an activity that is critical for mitochondrial physiology and muscle health.

## Introduction

Protein synthesis by mRNA translation is a highly coordinated process accomplished by the ribosome and an extensive array of eukaryotic initiation factors (eIFs) and additional ribosome-associated proteins. This machinery orchestrates many discrete steps which include the initiation of mRNA translation, translation elongation and termination as well as folding, targeting, and quality control of the nascent polypeptide (Joazeiro, 2019; Kramer et al., 2018; Pechmann et al., 2013). However, mechanisms coupling these discrete steps into a high-fidelity production line are just beginning to emerge.

Efforts over the past decades have primarily focused on deciphering mechanisms controlling canonical m7G cap-dependent mRNA translation initiation (Sonenberg and Hinnebusch, 2009). In canonical initiation, cap binding of eIF4E leads to the recruitment of eIF4G and eIF3 which, in turn, attracts the 43S pre-initiation complex consisting of the 40S ribosome loaded with methionyl-tRNA and additional eIFs to form the 48S initiation complex. The 48S complex then performs 5’-scanning of the mRNA to locate the start codon. Upon start codon recognition, the 60S ribosomal subunit joins with the 40S subunit to assemble an actively translating 80S ribosome, whereas eIFs are thought to be recycled.

While all eukaryotic mRNAs carry a m7G cap, only as few as ∼200 mRNAs appear to depend on the eIF4E cap binding protein (Hsieh et al., 2012; Morita et al., 2013; Thoreen et al., 2012; Truitt et al., 2015; Yanagiya et al., 2012), pointing to the existence of alternative non-canonical initiation pathways. Recent studies have implicated the 13-subunit eIF3 complex – the largest of the eukaryotic initiation factors - in several non-canonical modes of initiation (Wolf et al., 2019). Using PAR-CLIP, eIF3 was shown to regulate the translation of hundreds of mRNAs through interaction with structured mRNA motifs located in 5’-UTRs (Lee et al., 2015). Binding of eIF3 to one such motif present in the mRNA encoding c-Jun was shown to unmask a cryptic cap-binding activity of the eIF3d subunit, which is required for efficient cap-dependent but eIF4E-independent translation of c-Jun mRNA (Lee et al., 2016). eIF3 binding sites in mRNA frequently overlap with sites of N6 methyl adenosine (m6A) modification, and eIF3 was shown to bind to m6A either directly or through m6A reader and writer proteins (Choe et al., 2018; Meyer et al., 2015; Shi et al., 2017; Wang et al., 2015). Thus, eIF3 now appears to be involved in many different forms of canonical and non-canonical translation initiation, promoting ternary complex recruitment, binding of the 43S pre-initiation complex to mRNA, closed loop formation as well as scanning processivity and fidelity (Hinnebusch, 2017; Valášek et al., 2017; Wolf et al., 2019).

Although eIF3 has traditionally been thought to be released from ribosomes upon 40S-60S subunit joining in vitro (Peterson et al., 1979; Trachsel and Staehelin, 1979), recent evidence suggested that it can remain associated with 80S ribosomes during translation of short uORFs in order to facilitate re-initiation on downstream ORFs (Hronová et al., 2017; Mohammad et al., 2017). In addition, pull-down experiments demonstrated that yeast eIF3 forms stable in vivo interactions with factors involved in translation elongation (40S and 60S ribosomal proteins, eukaryotic elongation factors (eEFs), and tRNA synthetases), suggesting an unexplored role in integrating translation initiation with elongation (Sha et al., 2009).

By positional mapping of ribosomes at nucleotide resolution, we found that eIF3, through physical interaction with 80S ribosomes, promotes early translation elongation of select mRNAs encoding membrane functions. Our data suggest a model for a novel role of eIF3 in the coupling of translation initiation with elongation and co-translational membrane targeting, which is critical for mitochondrial physiology and muscle function.

## Results

### Cells deficient in eIF3e retain a partial eIF3 core complex

To explore the global role of eIF3 in mRNA translation at nucleotide resolution, we sought to perform ribosome profiling (Ingolia et al., 2009) in human cells upon knockdown (KD) of eIF3 subunit “e”. Knockdown of eIF3e in MCF-10A cells led to almost complete loss of eIF3e from high molecular weight eIF3 complexes resolved by native polyacrylamide gel electrophoresis (PAGE) (Fig. 1A). Similarly severe depletion was noted for eIF3d, a subunit known to directly interact with eIF3e and to be dependent of eIF3e for maintaining steady-state expression (Shah et al., 2016; Wagner et al., 2016; Yen and Chang, 2000). In addition, the vertebrate-specific subunit eIF3k as well as eIF3c were depleted, whereas core subunits eIF3a, eIF3b, and eIF3g were largely unaffected. A similar picture arose from quantitative LC-MS/MS analysis of sucrose density gradient fractions corresponding to 80S complexes of MCF-10A and HeLa cells, which showed that knockdown of eIF3e in these cell lines resulted in the depletion of the eIF3d/e module as well as eIF3k, eIF3c, and eIF3m (Fig. 1B). Notably, a roughly stochiometric residual eIF3 complex containing the subunits eIF3a, eIF3b, eIF3f, eIF3g, and eIF3i, which was previously described (Wagner et al., 2014), was clearly retained and appears to be joined to varying degrees by other eIF3 subunits present at reduced levels. Similar conclusions regarding the persistence of a residual eIF3 complex were reached by native PAGE and parallel biochemical purification and LC-MS/MS of eIF3 from MFC7 cells in which eIF3e was knocked down (Fig. S1A).

**Figure 1.**
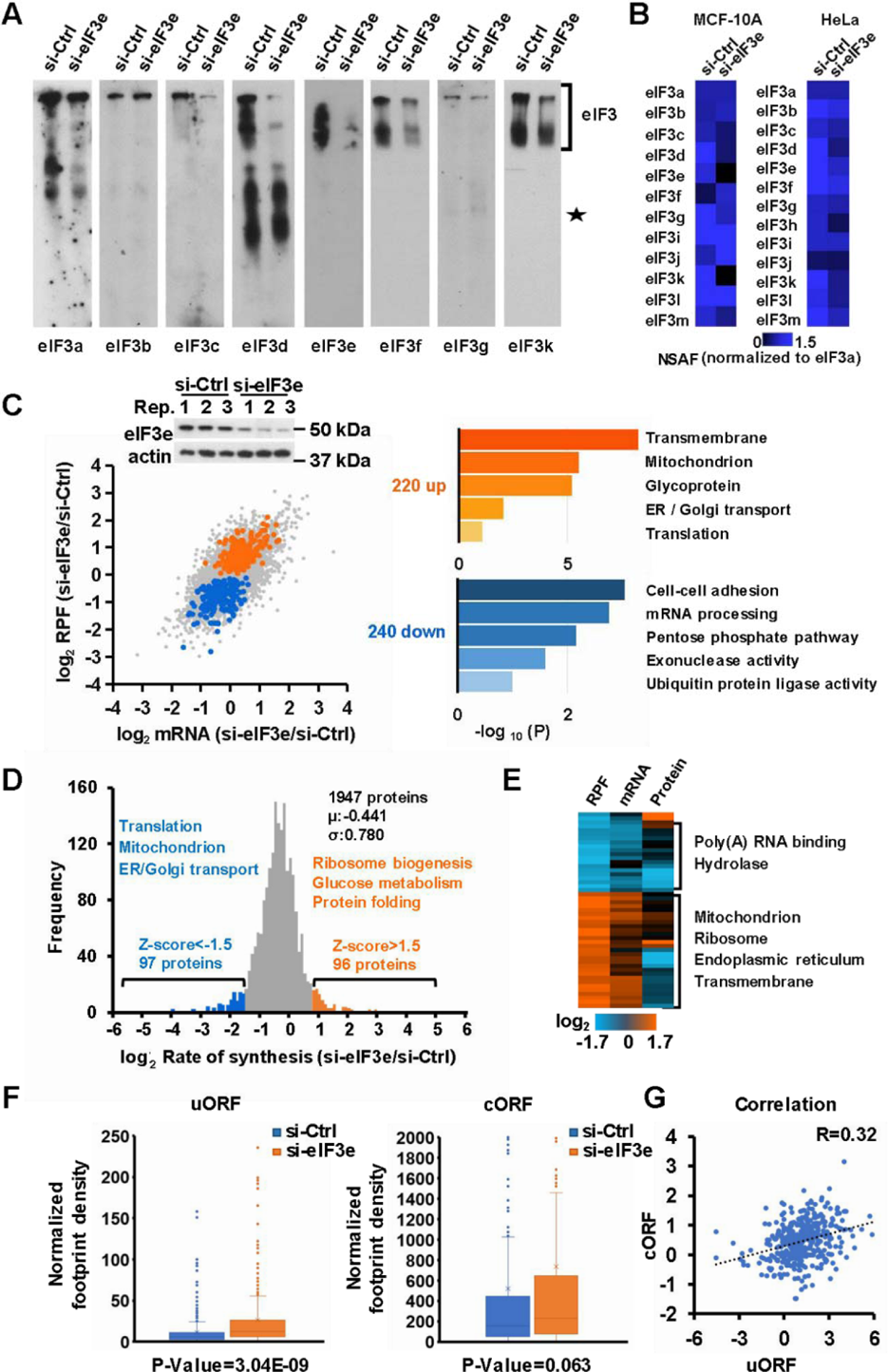
Effect of knockdown of eIF3e on ribosome density and protein synthesis. **A.** Cell lysate prepared from MCF-10A cells transfected with control (si-Ctrl) or eIF3e (si-eIF3e) small interfering RNAs was separated by native gel electrophoresis and blotted with the indicated antibodies directed against eIF3 subunits. The star denotes apparent non-specific bands reacting with the respective antibodies. **B.** Lysate of MCF-10A and HeLa cells transfected with control (si-Ctrl) or eIF3e (si-eIF3e) small interfering RNAs was separated by 10 – 50% sucrose density gradient centrifugation. The fraction corresponding to the 80S ribosomal peak was digested with trypsin and analyzed by LC-MS/MS. The heat map shows the relative quantification of eIF3 subunits based on normalized spectral abundance factors (NSAF). **C.** Ribosome profiling experiments were performed in triplicates with MCF-10A cells after knockdown of eIF3e for 72 hours. The method for analyzing changes in translational efficiency was adapted from Werner et al. (Werner et al., 2015). Average values of triplicate ribosome profiling (log2 RFP) and RNA-Seq (log2 mRNA) datasets were plotted in a scatter plot, and items changing with p < 0.01 in RFP but not mRNA were highlighted and subjected to functional term enrichment. For stringent analysis, the minimum number of reads was set to 10 and the p value cutoff was set to 0.01. Data points highlighted in orange correspond to 240 mRNAs with increased translational efficiency in eIF3e-KD cells, fitting the criteria log2 mRNA (si-eIF3e/si-Ctrl) p > 0.01, log2 RPF (si-eIF3e/si-Ctrl) p < 0.01, fold change > 0. The blue data points correspond to 220 mRNAs fitting the criteria: log2 mRNA (si-eIF3e/si-Ctrl) p > 0.01, log2 RPF (si-eIF3e/si-Ctrl) p < 0.01, fold change < 0. The right panel shows enriched functions in the up- and down-regulated groups of mRNAs. **D.** Protein synthesis as determined by pulsed SILAC. The graph highlights proteins whose synthesis is either down- or up-regulated upon knockdown of eIF3e. **E.** Heatmap comparing ribosome density (RPF) with mRNA abundance and protein synthesis (pSILAC). Log2 si-eIF3e/si-Control. **F.** Ribosome densities on 443 uORFs, their corresponding 5’-UTRs, and their subsequent cORFs. **G.** Correlation between ribosome densities on uORFs and cORFs.

### Loss of eIF3e curbs the synthesis of mitochondrial proteins despite paradoxical effects on ribosome occupancy

Highly reproducible ribosome profiling experiments performed with MFC-10A cells following eIF3e-KD (Fig. S2) revealed several hundred significant changes in translational efficiency (TE) defined as changes in ribosome protected fragments (RPF, p < 0.01) but not in total mRNA (p > 0.01) (Fig. 1C, Supplementary Data File 1). The group of 220 mRNAs with increased TE was enriched for those encoding mitochondrial and transmembrane proteins, in addition to proteins functioning in mRNA translation (Fig. 1C). Conversely, TE was decreased for a group of 240 mRNAs that was enriched in cell adhesion, mRNA processing and metabolic functions (Fig. 1C).

The increase in TE we observed for mRNAs encoding mitochondrial proteins in eIF3e-KD cells contradicted our previous demonstration in fission yeast that eIF3e promotes the synthesis of mitochondrial proteins (Shah et al., 2016). Consistent with the previous data, quantitative proteomics by pulsed SILAC (pSILAC) (Schwanhäusser et al., 2009) confirmed decreased synthesis of mitochondrial and transmembrane proteins in eIF3e-KD cells (Fig. 1D, Supplementary Data File 2). In contrast, the synthesis of proteins involved in glucose metabolism, ribosome biogenesis, tRNA aminoacylation, and protein folding was increased (Fig. 1D). However, most proteins encoded by mRNAs with increased TE for which pSILAC data were obtained (a total of 49) showed either reduced or largely unchanged synthesis (Fig. 1E, Supplementary Data File 3). Thus, eIF3e-KD results in reduced synthesis of mitochondrial and transmembrane proteins despite paradoxical increases in the apparent TEs of their encoding mRNAs.

As many as ∼50% of mammalian mRNAs are thought to contain upstream open reading frames (uORFs) that were proposed to impose negative control on the translation of downstream coding ORFs (cORFs) (Calvo et al., 2009; Johnstone et al., 2016). Since eIF3 is known to mediate re-initiation at uORFs thus affecting the translation of cORFs (Hronová et al., 2017; Szamecz et al., 2008), we explored the global impact of eIF3e on the translation of uORFs. We identified a total of 443 actively translated uORFs showing the characteristic 3 nucleotide periodicity in their ribosome density profiles (Supplementary Data File 4). We detected an increase in ribosome density on uORFs in eIF3e-KD cells (p = 3.04^-9^) (Fig. 1F). Curiously, this was reflected in a corresponding, though less pronounced, increase in ribosome density in the subsequent cORFs (p = 0.063) for an overall positive correlation between uORF and cORF ribosome density of 0.32 (Fig. 1G). This was also apparent at the level of individual mRNAs (Fig. S2C). Thus, while we did not see the inverse correlation between uORF and cORF ribosome occupancy described in previous reports (Calvo et al., 2009; Johnstone et al., 2016), our results are consistent with more recent studies finding a positive correlation (van Heesch et al., 2019; McGillivray et al., 2018). Regardless, these data strongly suggest that effects on uORF translation do not underlie the paradoxical relationship between ribosome occupancy of mRNAs and the synthesis of their encoded proteins we detected in eIF3e-KD cells.

### eIF3e is required for early translation elongation on a select class of mRNAs

In further attempts to reconcile the above paradox, we chose to compare the global patterns of ribosome distribution on mRNA of eIF3e replete and depleted cells. In a meta-analysis of normalized ribosome densities, we found a pronounced accumulation of ribosomes in a region between codons 25 and 75 in eIF3e-KD cells with a distinct peak at codon 50 (Fig. 3SA). No such accumulation was seen surrounding the stop codon or in the total RNA samples (Fig. S3A,B). From a total dataset of 19330 mRNAs, we identified 2683 mRNAs with a >2-fold increase in ribosome density between codons 25 and 75 (Fig. 2A, Fig. S3C, Supplementary Data File 5). This increase was reflected in a highly significant 5’ shift in global ribosome positioning along mRNAs (Fig. 2B). Selective ribosome accumulation in the region encompassing codon 50 was also apparent at the level of specific mRNAs (Fig. 2C, Fig. S3D). This data suggested that, due to abnormal, localized ribosome accumulation, global ribosome occupancy in eIF3e deficient cells is not a surrogate of protein synthesis and, in fact, shows an inverse relationship with apparent TE (Fig. 1E). Most mRNAs for which disruption of eIF3 increased early ribosome density were impaired in translational efficiency as indicated by reduced synthesis of their encoded proteins (Fig. 2D, Supplementary Data File 6). Thus, loss of eIF3 integrity causes a slowdown in early translation elongation resulting in localized ribosome accumulation between codons 25 – 75.

**Figure 2.**
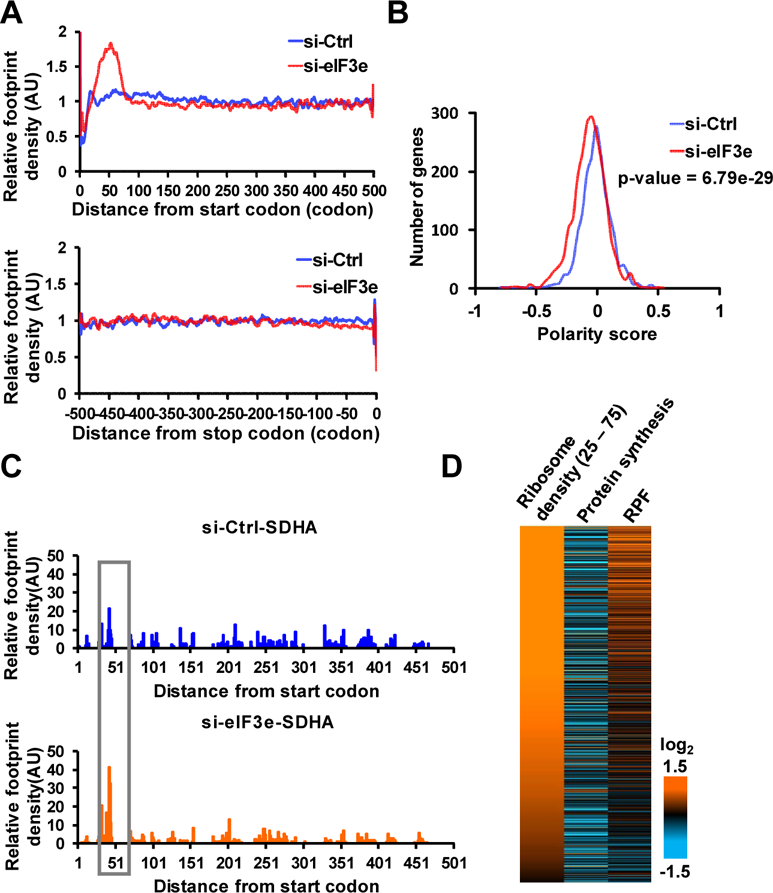
Effect of eIF3 on early translation elongation. **A.** Meta-analysis of 2683 mRNAs that show a pronounced accumulation of ribosomes in a region between codons 25 – 75. No such accumulation is seen at the stop codon. **B.** Global polarity score indicating the relative ribosome distribution towards the 5’ (negative score) or 3’ (positive score) region of mRNAs. **C.** Accumulation of ribosomes in the region between codons 25 and 75 in the mRNA encoding SDHA. **D.** Heatmap showing protein synthesis rates and total ribosome densities (RPF) of 1209 mRNAs with >2-fold increased ribosome density between codons 25 – 75.

### eIF3e-dependent mRNAs encode proteins with membrane functions and subunits of protein complexes sharing unique N-terminal features

We next sought to identify features that distinguish the group of 2683 mRNAs that depended on eIF3e for early elongation from eIF3e-independent mRNAs. The set of eIF3e-dependent mRNAs showed strong enrichment of functional pathways which were dominated by RNA metabolism, protein homeostasis, and membrane-associated processes (Fig. 3A, Supplementary Data File 7). This encompassed 1229 membrane proteins, hundreds of mitochondrial, endosomal, lysosomal and secreted proteins as well as subunits of large protein complexes, including the ribosome, the spliceosome, the proteasome, and the nuclear pore complex (Fig. 3B). As a group, these proteins have increased hydrophobicity and positive charge within the N-terminal ∼25 residues (Fig. 3C,D), which are well-established characteristics of mitochondrial targeting sequences (Omura, 1998).

**Figure 3.**
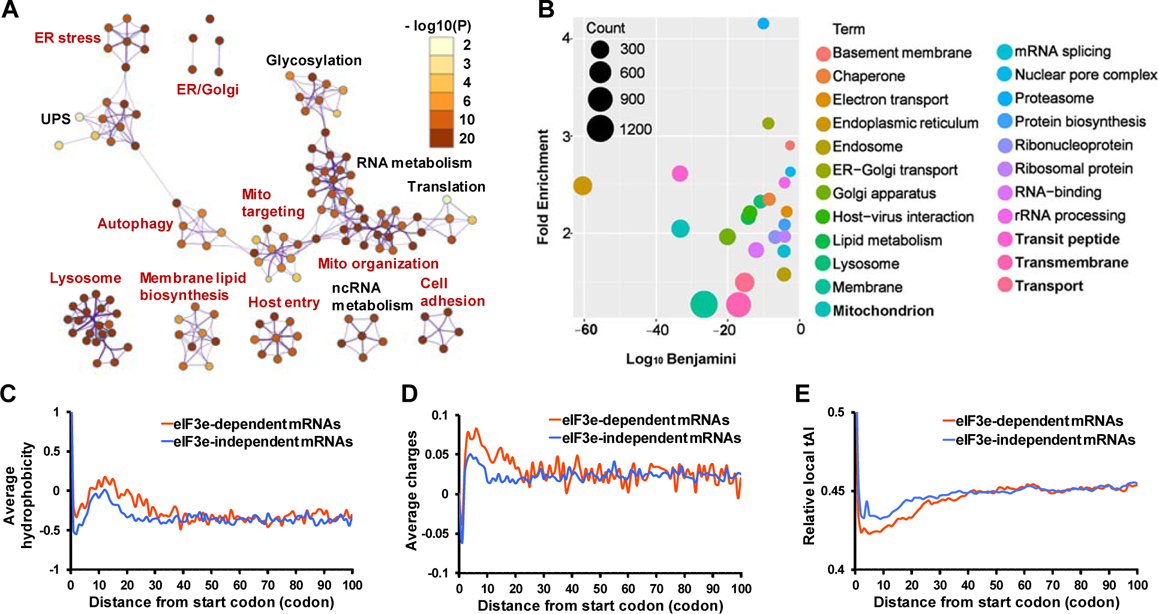
Features of proteins encoded by eIF3e-dependent mRNAs. **A.** Network of Gene Ontology terms enriched in the set of 2683 mRNAs with increased ribosome density between codons 25 and 75. Clusters denoted in red indicate membrane-associated functions. UPS, ubiquitin-proteasome system. **B.** Enrichment of Uniprot Keywords in the set of 2683 mRNAs with increased ribosome density between codons 25 and 75. The graph plots the fold enrichment of functional Uniprot terms against the significance of enrichment. The size of the circles represents the relative number of mRNAs in each category. **C.** Average hydrophobicity of the proteins encoded by the 2683 mRNAs that show high ribosome density in eIF3e depleted cells (“eIF3e-dependent mRNAs”) versus the proteins encoded by the remaining set of 16229 mRNAs that do not show the elongation block (“eIF3e-independent mRNAs”). **D.** Average positive charge (lysine and arginine residues) of the protein sets described in c). **E.** Relative local tRNA adaptation index (tAI) of the protein sets described in C.

We did not detect any common RNA or amino acid sequence motifs apparent in the group of 2683 mRNAs that might underly their dependence on eIF3 for early translation elongation. Notably, there was no obvious difference in the Kozak consensus context (Kozak, 1986) between eIF3-dependent and -independent mRNA sets (Fig. S3E). The observation that optimality of the nucleotide context surrounding the start codon does not underlie eIF3e dependency is consistent with eIF3e promoting translation elongation rather than initiation on this set of mRNAs. Depletion of tRNAs may impede elongation, but we likewise found no difference in tRNA levels between eIF3e-KD and control cells (Fig. S3F). However, eIF3-dependent mRNAs had a lower tRNA adaptation index in the region encoding the N-terminal ∼40 amino acids (Fig. 3E). The prevalence of rare codons suggests that eIF3-dependent mRNAs may begin their translation at a slower pace than eIF3-independent mRNAs, a feature known to be important for efficient synthesis of membrane proteins (Acosta-Sampson et al., 2017).

### eIF3 interacts with 80S ribosomes during translation of the first ∼60 codons

To mechanistically address the function of eIF3 in elongation, we asked whether the initiation factor physically interacts with elongating 80S ribosomes. On sucrose density gradients, eIF3 subunits were strongly enriched in 40S monosomal fractions but also co-eluted with polysomes (Fig. 4A), suggesting association with actively translating 80S ribosomes. A similar pattern was seen for the cap binding protein eIF4E (Fig. 4A). Unlike the ribosomal proteins RPL7 and RPS19, eIF3 subunits and eIF4E did not increase proportionally with the density of the polysomal fractions (Fig. 4A), indicating that initiation factors might associate with only a subset of 80S ribosomes.

**Figure 4.**
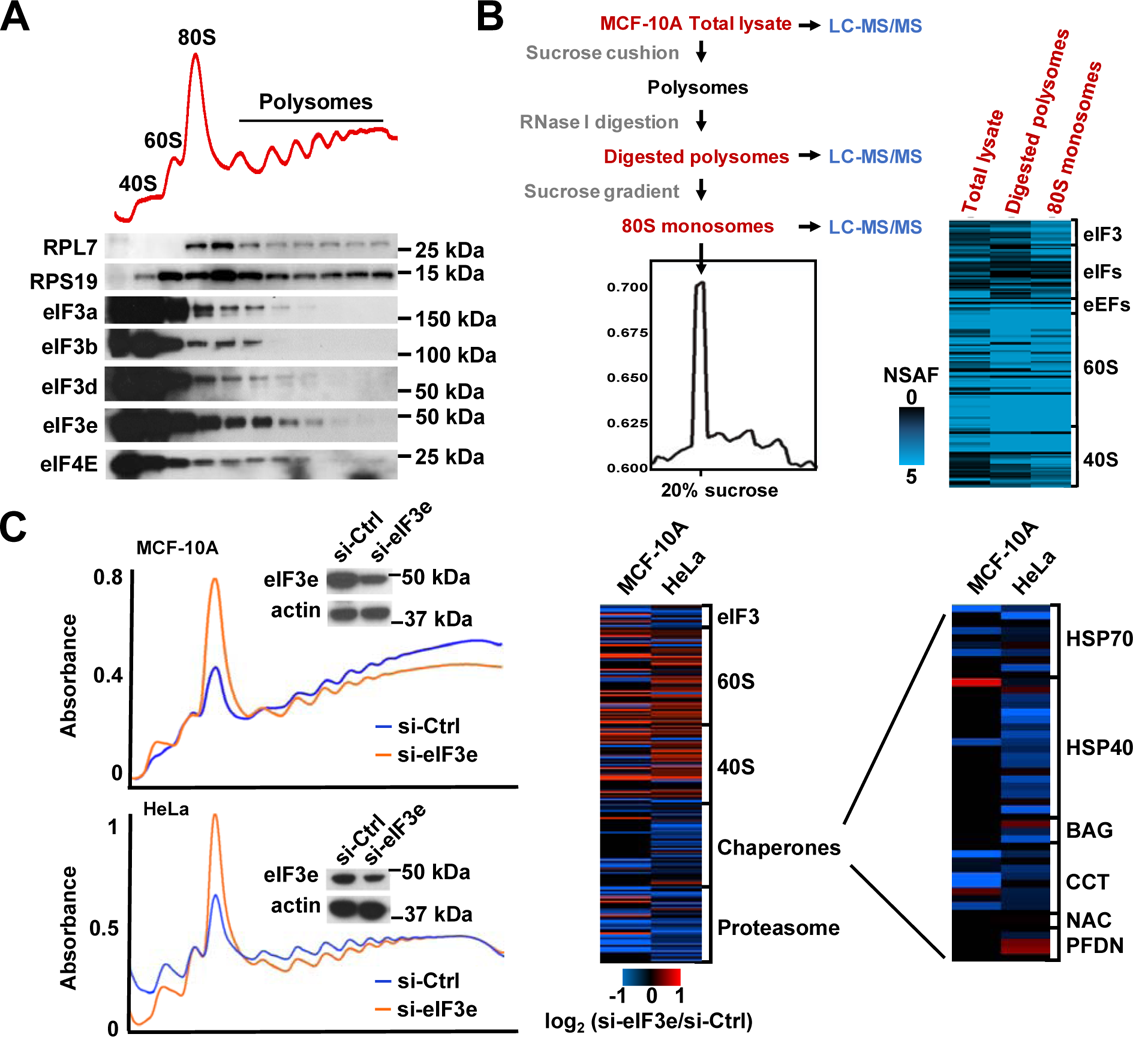
eIF3 associates with 80S ribosomes and protein quality control factors. **A.** Polysome profiling of MCF-10A cells by sucrose density gradient centrifugation. The elution profiles of the indicated translation proteins were determined by immunoblotting. **B.** Purification scheme and heatmap showing the abundance of the indicated protein groups in total MCF-10A cell lysate, RNAse I digested polysomes, and purified 80S monosomes as determined by quantitative LC-MS/MS. **C.** Lysate of MCF-10A and HeLa cells 72 h after knockdown of eIF3e were separated on sucrose density gradients, followed by quantitative LC-MS/MS of 80S monosome fractions. The heatmap shows Log2 changes between si-Control and si-eIF3e cells.

To exclude that eIF3 co-elution with polysomes was due to the presence of scanning 43S pre-initiation complexes on polysomal mRNAs, we digested polysomal fractions with RNAse I and purified the resulting 80S ribosomal complexes by sucrose density gradient fractionation. LC-MS/MS revealed copious amounts of eIF3 in addition to 40S and 60S ribosomal proteins in purified 80S complexes (Fig. 4B, Supplementary Data File 8). Purified 80S complexes also contained other proteins we had previously found to interact with eIF3, including initiation factors (eIF4E, eIF4G), elongation factors (eEFs), and the 26S proteasome (Sha et al., 2009). Quantitative proteomics of 80S fractions obtained upon knockdown of eIF3e in two different cell lines, MCF-10A and HeLa, revealed a decrease in the abundance of protein quality control factors, including HSP70 and HSP40 chaperones, the CCT/TRiC chaperonin, and the proteasome despite increased amounts of ribosomal proteins (Fig. 4C, Supplementary Data File 9). This depletion suggests that eIF3, via its eIF3e subunit, serves to recruit protein quality control factors to 80S ribosomes, possibly in order to promote translation elongation.

To map the location of eIF3-associated 80S ribosomes on mRNA, we performed selective ribosome profiling under native conditions without cross-linking reagents (Becker et al., 2013). Following digestion of MCF-10A cell lysate with RNAse I, eIF3-associated ribosomes were immunopurified using eIF3b or eIF3e antibodies which efficiently precipitate the eIF3 complex (Fig. S1B), and ribosome protected fragments (RPFs) were extracted from the immunopurified eIF3-80S complexes for sequencing (Fig. S4A). Enrichment analysis of the eIF3-bound RPFs relative to total RPFs was used to map the presence of eIF3-associated 80S ribosomes on mRNA. The analysis of eIF3b and eIF3e selective ribosome profiling datasets revealed a consistent enrichment of eIF3-80S complexes within the first ∼60 codons on the group of 2683 mRNAs identified by ribosome profiling as being dependent on eIF3e for efficient translation elongation (Fig. 5A). An unbiased search for mRNAs displaying enrichment of eIF3 on 80S ribosomes in the 5’ part of the coding region identified classes of 2543 (eIF3b) and 4623 (eIF3e) transcripts, respectively (Fig. 5B, Supplementary Data File 10), which showed a highly significant overlap with the set of 2683 eIF3-dependent mRNAs (P < 1.25^-49^, Fig. S4B). Both mRNA sets were enriched for the same functional categories, dominated by mitochondrial proteins and membrane-associated processes (Fig. 5C, Supplementary Data File 11). Taken together, these findings suggest that eIF3 promotes early translation elongation through association with translating 80S ribosomes, presumably serving to recruit protein quality control factors.

**Figure 5.**
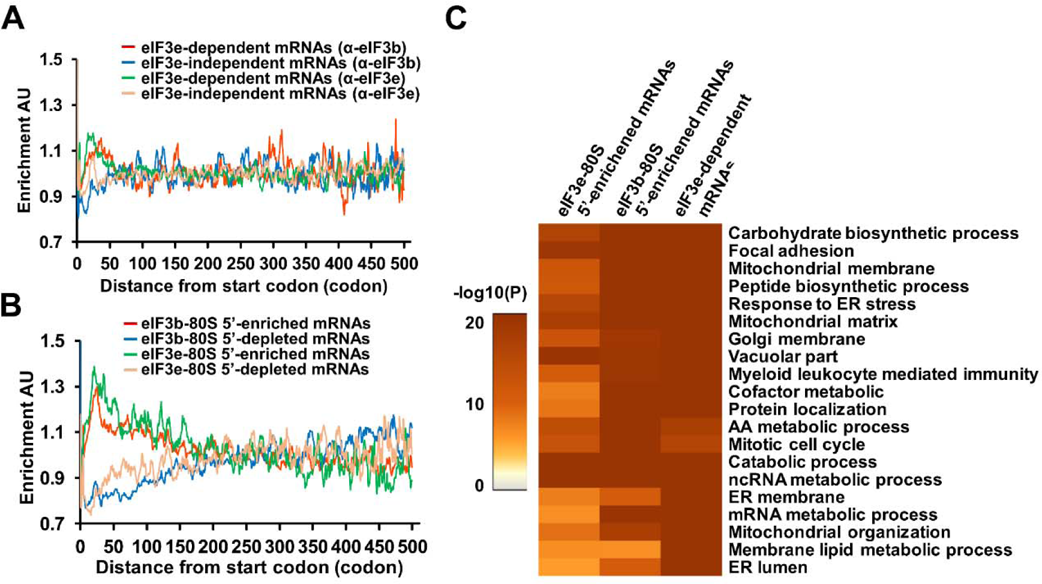
eIF3 associates with 80S ribosomes early during translation. **A.** Selective ribosome profiling of 80S-eIF3 interactions using α-eIF3b and α-eIF3e antibodies. Cumulative distribution of eIF3-associated 80S ribosomes on the set of 2683 eIF3e-dependent mRNAs compared to the set of 16229 eIF3e-independent mRNAs as established by selective profiling of eIF3b- and eIF3e-associated ribosomes. **A. B.** Distribution of eIF3b-associated 80S ribosomes on a set of 2543 mRNAs showing global 5’ accumulation of eIF3b-associated ribosomes relative to all ribosomes (red graph, polarity difference <0, “eIF3-80S 5’-enriched mRNAs”, see Methods for details). The blue graph represents a reference set of 5204 mRNA with a polarity difference >0 (“eIF3-80S 5’-depleted mRNAs). The green and pink graphs show the same data for eIF3e (4623 mRNAs with polarity difference <0, 4179 mRNAs with polarity difference >0). **B.** Overlap in the enrichment of functional terms in the set of 2683 eIF3-dependent mRNAs and the sets of mRNAs with increased density of eIF3b- or eIF3e-associated ribosomes towards the 5’ end (“eIF3-80S 5’-enriched mRNAs”).

### eIF3e is required for mitochondrial physiology and skeletal muscle health

To determine whether the mRNA selective requirement of eIF3e for protein synthesis at the level of translation elongation uncovered here gives rise to corresponding cellular phenotypes, we assessed the ultrastructural morphology of eIF3e-KD cells by transmission electron microscopy. Reflecting the functional pathways enriched in eIF3-dependent and eIF3-bound mRNA sets, eIF3e-KD cells showed a marked disturbance in organelle structure with accumulation of electron sparse vesicles that were filled with multilamellar membranous contents, morphological features consistent with lysosomes (Fig. 6A). To decipher the identity of the vesicles accumulating in eIF3e-KD cells, we used transfection of a mCherry-EGFP-LC3B fusion protein as an autophagosomal/lysosomal marker. In si-Control cells, mCherry single positive and mCherry-EGFP double positive yellow vesicles coexisted at relatively low numbers, indicating intact autophagosomal and lysosomal compartments (Fig. 6B, see Fig. S5D for additional examples). Upon the addition of the lysosomal inhibitor bafilomycin A1, more yellow puncta were observed, a finding that is consistent with inhibition of lysosomal degradation at constant autophagic flux (Fig. 6B, Fig. S5D). In eIF3e-KD cells, the majority of puncta appeared red, but yellow puncta were observed upon treatment with bafilomycin A. This indicates intact autophagic flux but increased lysosomal load in eIF3e-depleted cells, possibly to clear defective mitochondria. Supporting this idea, mitochondrial cristae appeared dilated with fuzzy borders in eIF3e-KD cells (Fig. 6A). Consistent with mitochondrial dysfunction, metabolic flux analysis revealed a decrease in mitochondrial oxygen consumption rate that depended on the dose of eIF3e siRNA (Fig. 6C, Fig. S5E).

**Figure 6.**
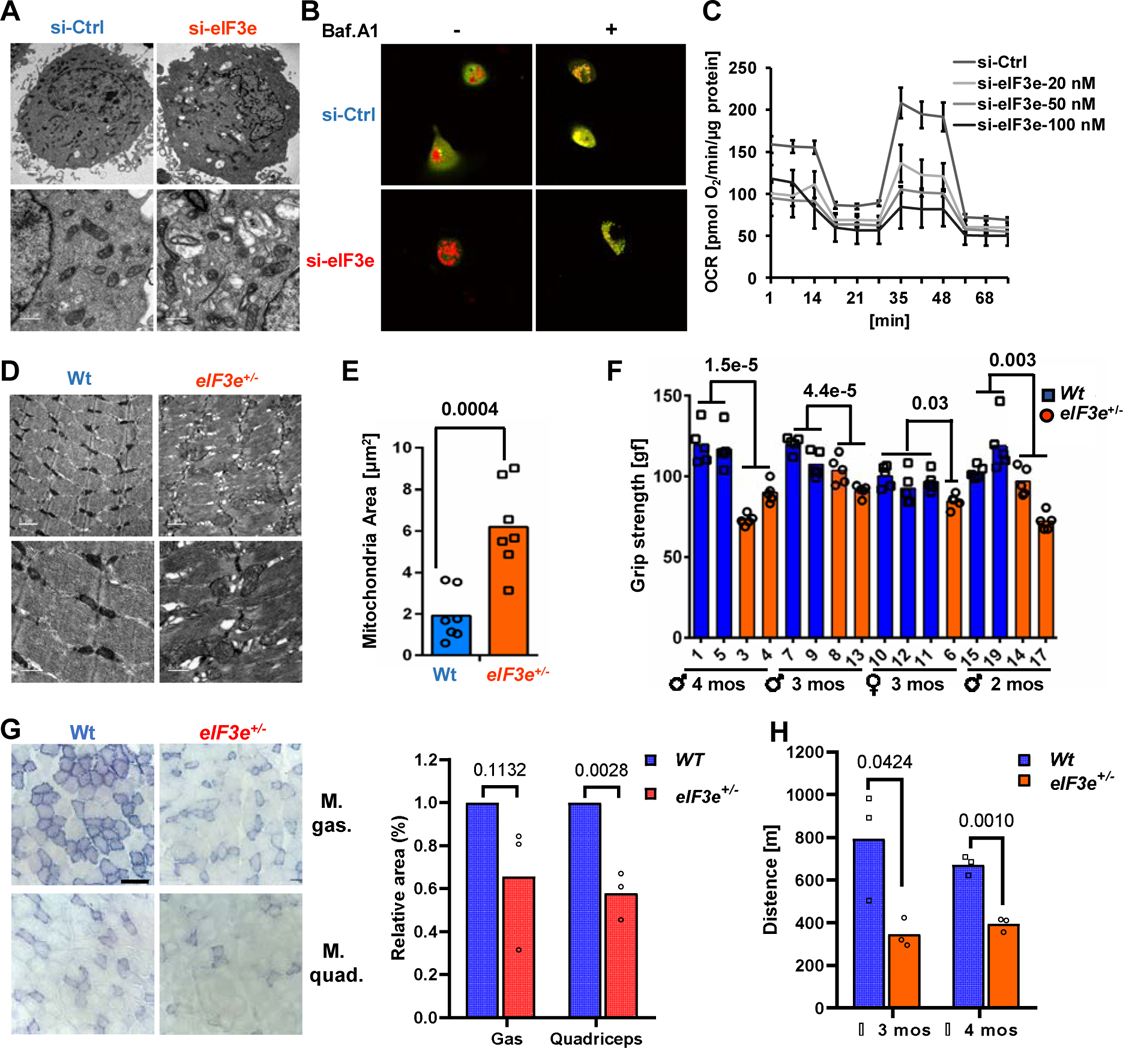
Effect of eIF3e on cell a d muscle physiology. **A.** Transmission electron microg hs on MCF-10A cells upon knockdown of eIF3e for 72 hours. See Fig. S5A for eIF3e knockdown efficiency. Size bars: 2 μm (top panel), 0.5 μm (bottom panel). **B.** Live cell imaging of MCF-10A cells transfected with a plasmid driving mCherry-EGFP-LC3B fusion protein. Lysosomes in red, autophagosomes in yellow. See Fig. S5B for eIF3e knockdown efficiency. **C.** Mitochondrial oxygen consumption rate (OCR) of MCF-10A cells exposed to increasing concentrations of si-eIF3e for 72 hours. OCR was measured in a FX Flux Analyzer. See Supplementary Fig. 5C for eIF3e knockdown efficiency. **D.** Electron micrographs of hindlimb skeletal muscle from wildtype and eIF3^+/-^ heterozygous knockout mice. See Fig. S6 for genotyping. Size bars: 1 μm (top panel), 0.5 μm (bottom panel). **A. E.** Quantification of mitochondrial area in electron micrographs of skeletal muscle from wildtype (n = 6) and eIF3^+/-^ (n = 6) mice. **B. F.** Grip test to measure muscle strength in wildtype and eIF3^+/-^ heterozygous knockout mice. **C. G.** NADH:ubiquinone oxidoreductase (complex I) activity in gastrocnemius and quadriceps muscles of wildtype and eIF3^+/-^ mice. Size bar: 100 μm. The bar graph to the right shows a quantification of the relative area of positive complex I activity staining in muscle of three independent animals. **D. H.** Mice were run at 10 m/min for 10 min, and speed was increased by 2 m/min every 1 min until mice were exhausted. Distance run until the mice failed to climb the treadmill within 5 s despite electric stimulation was recorded and plotted.

To corroborate the function of eIF3e in mitochondrial physiology at the organismal level, we generated eIF3e knockout mice. Unlike homozygous eIF3e knockout mice which were not obtained and thus appear inviable (Fig. S6A,B), eIF3e^+/-^ knockout mice show no overt phenotype except for a ∼10% reduction in body weight (Fig. S6C). A phenotype indicative of mitochondrial dysfunction was, however, observed in skeletal muscle of heterozygous eIF3e^+/-^ knockout mice we created. Muscle of 10 to 18 week old eIF3e^+/-^ mice showed a pronounced accumulation of hyperfused intramyofibrillar mitochondria (Fig. 6D,E, Fig. S6D), possibly an adaptive response to energy stress caused by eIF3e deficiency (Tondera et al., 2009). In addition, severe ultrastructural damage to the sarcomers with irregular and diffuse Z-disks and degeneration of the contractile elements was observed (Fig. 6D, Fig. S6D). Significant impairment of neuromuscular function was indicated by a progressive reduction in grip strength of male and female eIF3e^+/-^ mice (Fig. 6F).

To directly assess mitochondrial function in skeletal muscle of eIF3e+/-mice, we measured NADH:ubiquinone oxidoreductase (Complex I) activity in frozen sections using the NADH-tetrazolium reductase reaction. Notably, 16 of 38 nuclear encoded CI subunits, including most subunits of the catalytic matrix arm, are within the set of 2683 mRNAs requiring eIF3e for efficient translation elongation (Supplementary Data File 5). We observed strongly diminished Complex I activity in type II fast twitch quadriceps and gastrocnemius muscles (Fig. 6G). Consistent with this decrease, eIF3e+/-mice showed lower endurance in uphill treadmill running (Fig. 6H). Taken together, these findings suggest that the translation elongation function of eIF3e is required for mitochondrial physiology and skeletal muscle health.

## Discussion

### mRNA selective promotion of translation elongation via eIF3-80S interaction

Mapping the positioning of ribosomes on mRNA, we established an unanticipated role of eIF3 in mRNA selective early translation elongation. Our studies suggest that eIF3 engages with ribosomes in at least two different fashions (Figure 7): 80S ribosomes translating mRNAs encoding soluble cytoplasmic and nuclear proteins loose eIF3 within the first few codons. A second set of mRNAs encoding membrane-associated functions retains eIF3 on the 80S ribosome for up to ∼60 codons. Retention of eIF3 on translating 80S ribosomes was recently also observed by chemical cross-linking (Bohlen et al., 2019; Wagner et al., 2019). Cryo-EM studies demonstrated that eIF3 undergoes conformational changes during the initiation reaction (Eliseev et al., 2018; Simonetti et al., 2016) that are consistent with persistence on the 80S ribosome. During scanning, eIF3b – and presumably other subunits of the yeast-like core complex (eIF3i and eIF3g) - move from the solvent exposed side of the 40S ribosome to the intersubunit side, thus preventing subunit joining. Upon start codon recognition, however, eIF3b (and eIF3i, eIF3g) relocate back to the solvent exposed side of the 40S ribosome, a conformational change accommodating 60S subunit joining without release of eIF3 (Eliseev et al., 2018; Simonetti et al., 2016).

**Figure 7.**
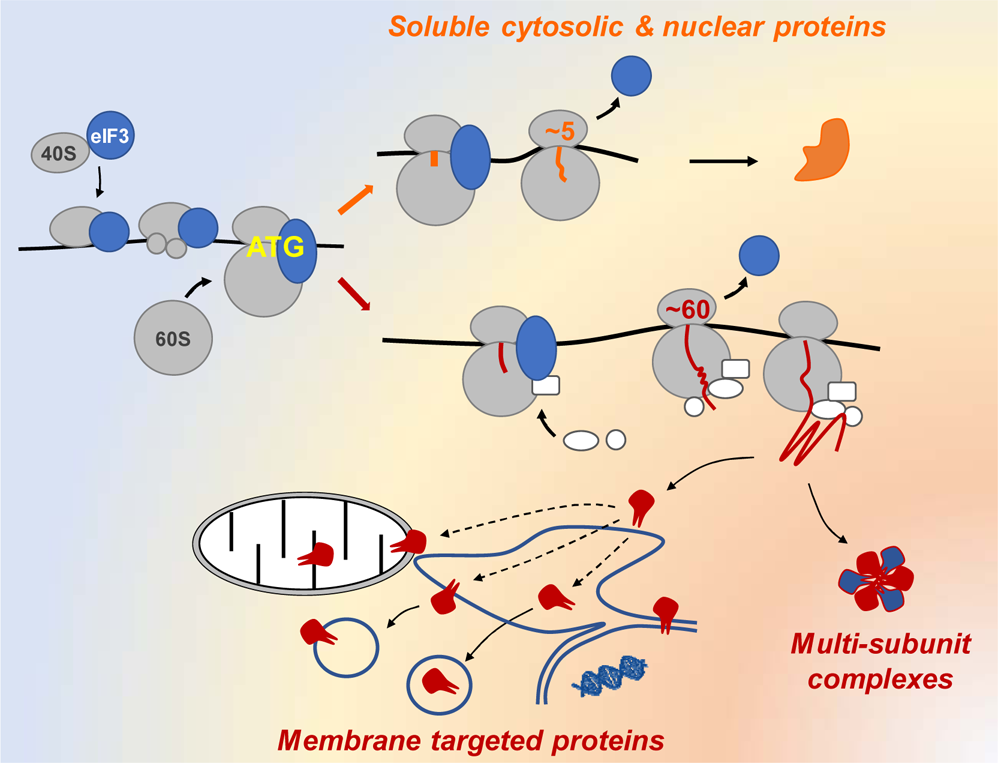
Model for the role of eIF3 in translation initiation, elongation, and targeting of nascent proteins to membranes and into protein complexes.

Our data further suggest that upon knockdown of eIF3e, a residual complex containing roughly stoichiometric amounts of subunits eIF3a, eIF3b, eIF3f, eIF3g, and eIF3i is retained, which appears sufficient for accomplishing the task of initiation. This raises the intriguing possibility that a sub-complex akin to the 5-subunit core complex found in *S. cerevisiae* may be proficient in initiation in mammalian cells. Remaining subunits may equip eIF3 with additional functionality in a modular fashion with the eIF3d-eIF3e module mediating early translation elongation. This modular architecture may enable eIF3 multi-functionality in integrating translation initiation with elongation and protein quality control.

### Potential mechanisms involved in eIF3e-mediated translation elongation

How might eIF3 promote translation elongation? We find it unlikely that eIF3 plays a role akin to eIF5A, which is a general promoter of elongation at the level of peptide bond formation at problematic motifs containing proline and aspartate (Schuller et al., 2017). Our ribosome profiling data did not provide evidence for ribosomal pausing at specific motifs. Rather, ribosomes lacking eIF3 are pausing in a defined region between codons 25 and 75 which approximately coincides with the emergence of nascent chains from the ribosome exit tunnel. This invokes a model (Figure 7) according to which, for a select group of mRNAs, eIF3 recruits factors to the 80S ribosome which receive the nascent chains to target them to their subcellular destinations, including the endoplasmic reticulum membrane for further sorting into the endosomal and secretory pathways as well as the mitochondrial retrieval pathway (Hansen et al., 2018). In a similar fashion, factors recruited by eIF3 may facilitate the co-translational formation of protein-protein interactions (Duncan and Mata, 2011; Shiber et al., 2018) required for the assembly of large protein complexes such as the proteasome, actin filaments, or, indeed eIF3 itself (Bohlen et al., 2019; Wagner et al., 2019). Since eIF3-dependent mRNAs are enriched for rare codons downstream of the AUG, slow initial progression of 80S ribosomes may facilitate eIF3-mediated factor recruitment.

Our quantitative 80S proteomics indicates that eIF3 serves to recruit HSP70 chaperones, including the ribosome-associated complex (RAC; HSPA14), and the CCT/TRiC chaperonin to early translating ribosomes. Both groups of chaperones are known to assist in co-translational protein folding (Albanèse et al., 2006; McCallum et al., 2000) as well as in the assembly of multi-subunit protein complexes (Melville et al., 2003). CCT/TRiC promotes the synthesis of membrane proteins, including some of those encoded by mRNAs contained within our list of 2683 mRNAs depending on eIF3e for early elongation (Génier et al., 2016). Likewise, budding yeast SSB-type HSP70 proteins mediate efficient synthesis of membrane and mitochondrial proteins (Döring et al., 2017). Unlike with budding yeast SSB-type HSP70s which are anchored to the ribosome, it remains unclear how mammalian HSP70s and CCT/TRiC are recruited to nascent chains. Our results raise the intriguing possibility that 80S-bound eIF3 mediates this recruitment within a particle we have previously named the “translasome” (Sha et al., 2009).

HSP70 and CCT/TRiC may serve as “holdases” to keep the nascent chains unfolded until they have been properly targeted to membranes or into interactions with their binding partners. If the recruitment is inefficient due to lack of eIF3, ribosomes may pause until the nascent chains can be properly directed. Alternatively, eIF3-recruited factors may actively promote elongation by exerting a mechanical force on the nascent chain which is relayed to the peptidyl transferase center (Goldman et al., 2015; Waudby et al., 2019). For example, mechanical pulling by the bacterial Sec translocon can promote translation elongation of translocating proteins (Goldman et al., 2015). Therefore, failure to recruit factors to the ribosome that promote protein folding and targeting to ER and mitochondrial translocons may reduce translation kinetics upon emergence of the nascent chains from ribosomes.

An intriguing question remains how mRNAs are selected to retain eIF3 on the 80S ribosome. It appears that this decision is made while the nascent peptide is still deep within the ribosome exit tunnel. Similar to mechanisms thought to govern SRP pre-recruitment to ribosomes even before the emergence of signal sequences (Chartron et al., 2016), the trigger for early loss versus prolonged retention of eIF3 may be hard-wired in the mRNAs. Notably, the list of eIF3e-dependent mRNAs shows a highly significant overlap (P < 7.181^-74^, Fig. S7A) with a previously identified list of mRNAs that bind eIF3a in their 5’-UTRs (Meyer et al., 2015). Both lists also enrich the same functional categories dominated by membrane-associated processes (Fig. S7B, Supplementary Data File 12). This suggests the possibility that direct eIF3-mRNA interactions select those mRNAs which will efficiently retain eIF3 during early elongation. It is also possible that early translation speed contributes to eIF3 retention, with mRNAs having suboptimal early codons retaining eIF3 longer.

### Role of eIF3e in mitochondrial and muscle physiology

Consistent with its mRNA selective elongation function, loss of eIF3 leads to decreased synthesis of mitochondrial proteins, a deficit in respiration, increased lysosomal load, decreased complex I activity, and apparent hyperfusion of mitochondria in skeletal muscle. In addition, sarcomeric structure is severely disturbed with muscle fiber disruption and irregular Z-disks. This finding is consistent with the established role of CCT/TRiC in the synthesis and folding of actin and myosin (Frydman, 2001). Indeed, CCT/TRiC was recently shown to be required for muscle actin folding and sarcomere assembly (Berger et al., 2018). It is thus conceivable that the skeletal muscle phenotype observed in eIF3e^+/-^ mice is, at least in part, due to inefficient recruitment of CCT/TRiC to 80S ribosomes. Many actin and myosin encoding mRNAs are on the lists of eIF3-regulated mRNAs we identified (Supplementary Data File 5 and 10). This list also includes a series of sarcomeric and Z-disk proteins (CAPZB, CD81, CFL1, CFL2, CRP1, CRP2, MESDC2, MYH10, MYOF, PKP2, TPM4, VCL). The combined demands for high mitochondrial energy production as well as precisely targeted assembly of complex sarcomeric protein structures may render skeletal muscle particularly dependent on eIF3’s function in translation elongation, protein targeting, and quality control.

## Acknowledgments

We thank E.-L. Eskelinen for evaluation of electron micrographs, M. Hansen for the pDest-mCherry-EGFP-LC3B plasmid, C. Hu for graphics support, and all members of the Wolf and Yang labs for discussion. The National Center for Protein Science (Beijing) is thanked for MS services. Support of the Platforms of Genome Sequencing and High-Performance Computing of the National Protein Science Facility (Beijing) at Tsinghua University is thankfully acknowledged. D.A.W.’s lab at Xiamen University is funded through grant 81773771 from the National Science Foundation of China and the 1000 Talent Program. During the initial phase of this work, D.A.W. received partial salary support from the US National Institutes of Health (grants R21 CA190588, R01 GM105802, and R01 GM121834). X. Y.’s lab is funded by the National Key Research and Development Program, Precision Medicine Project (2016YFC0906001), the National Natural Science Foundation of China (91540109 and 31671381) and the 1000 Talent Program (Youth Category).

## Author contributions

Conceptualization, Y.L. and D.A.W.; Methodology, Y.L., L.H., H.D., G.T., F.L. and X.X.; Formal Analysis, F.L., X.X., X.Y., Y.L.; Investigation, Y.L., F.L., L.H., H.D., L.S., J.F., X.X., G.T.; Writing – Original Draft, D.A.W.; Writing – Review & Editing, Y.L., F.L., L.H., L.S., J.F., H.D., X.X., G.T., Y.C., X.Y., D.A.W.; Visualization, Y.L., F.L. X.Y., D.A.W.; Supervision, Y.C., X.Y., and D.A.W.; Funding Acquisition, Y.C., X.Y., and D.A.W.

## Author information

Data are deposited in Gene Expression Omnibus (GSE131074) and Peptide Atlas (PASS01378). The authors declare no competing interests.

## Key Resources Table

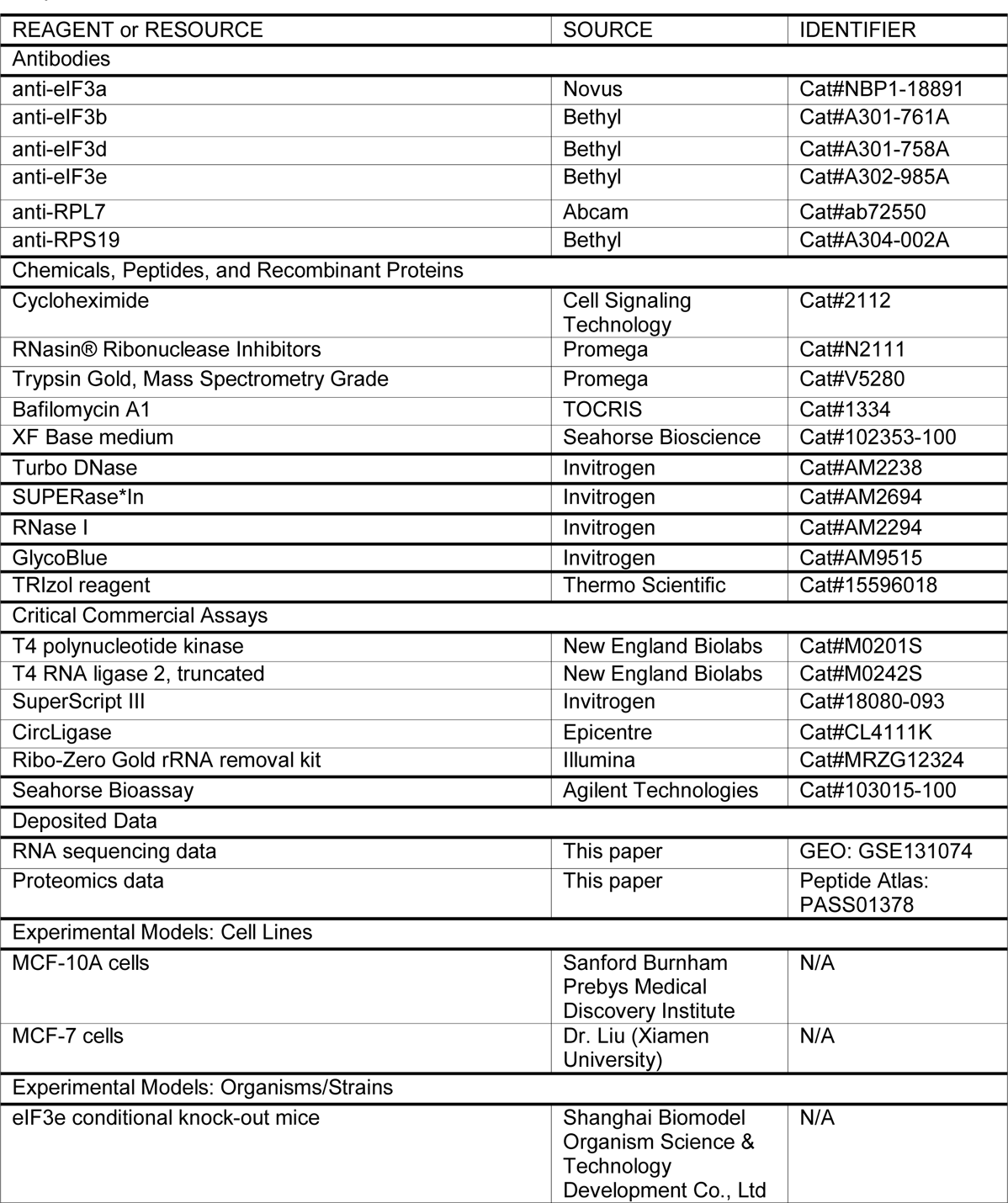
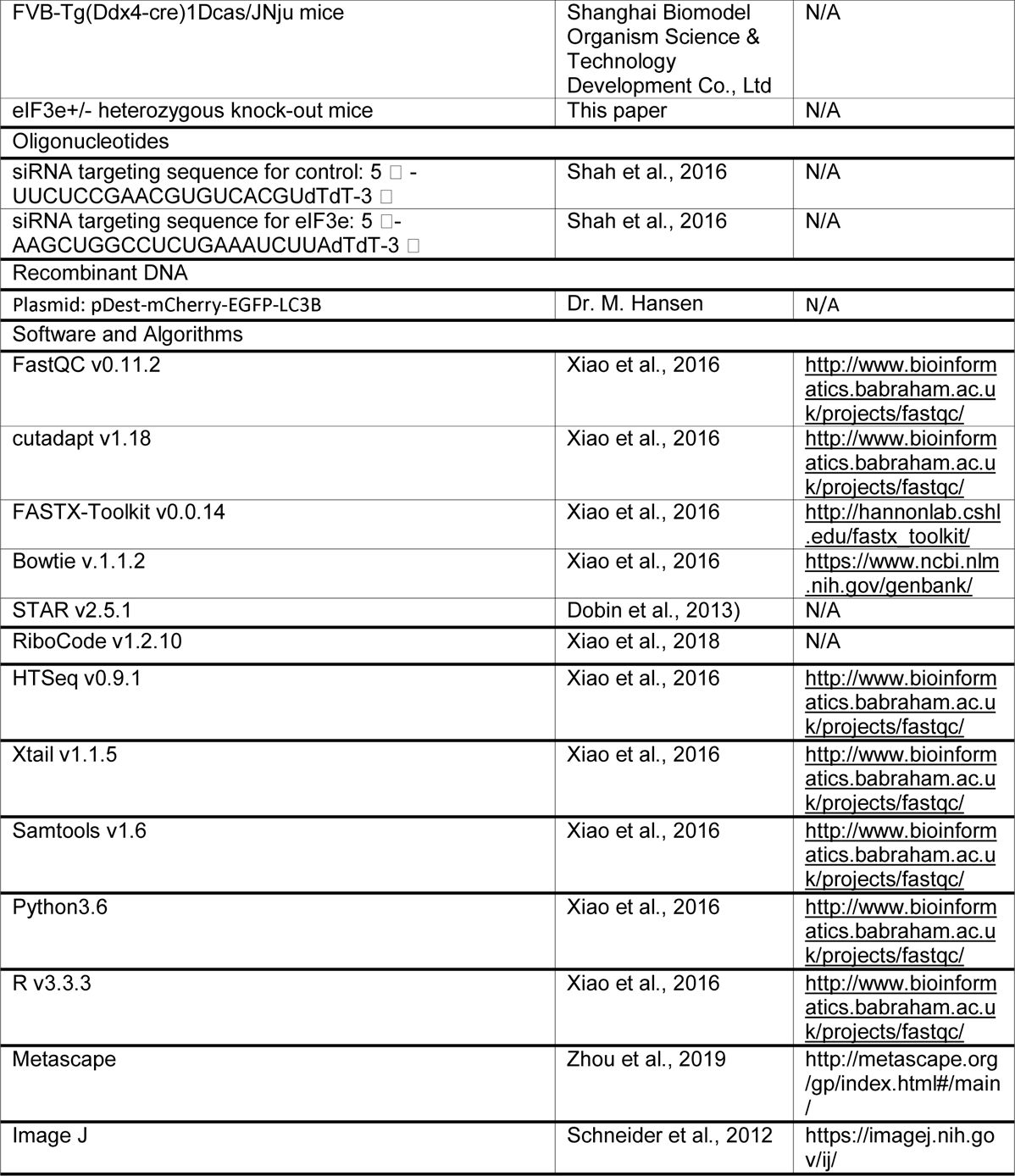

## Contact for Reagent and Resource Sharing

Further information and requests for reagents may be directed to, and will be fulfilled by the corresponding author Dieter A Wolf

## Method Details

### Tissue culture

MCF-10A cells were cultured in complete growth medium consisting of 50% Dulbecco’s modified Eagle’s medium and 50% Ham’s F12 medium supplemented with 5% fetal bovine serum, 20 ng/mL epidermal growth factor, 10 μg/mL insulin, 5 μg/mL hydrocortisone, 1 unit/mL penicillin, and 1 μg/mL streptomycin under a humified environment with 5% CO_2_ at 37°C. MCF-7 and HeLa cells were cultured in complete growth medium consisting of Dulbecco’s modified Eagle’s medium supplemented with 10% fetal bovine serum, 1 unit/mL penicillin, and 1 μg/mL streptomycin under a humified environment with 5% CO_2_ at 37°C. Cells were authenticated by short tandem repeat sequencing and determined to be free of mycoplasma.

### Knockdown of eIF3e

1 × 10^7^ MCF-10A cells were seeded and grown for 24 hours to a density of ∼40%. 10 M control or eIF3e si-RNA were transfected using Lipofectamine 2000 transfection reagent for 72 h before harvesting cells for protein or RNA preparations. The si-RNA oligo sequences were: si-control: 5 L -UUCUCCGAACGUGUCACGUdTdT-3 L; si-eIF3e: 5 L-AAGCUGGCCUCUGAAAUCUUAdTdT-3 L (Shah et al., 2016).

### Western blot

Cells were scraped in 0.1 mL SDS sample buffer (60 mM Tris/Cl, pH 6.8, 5% beta-mercaptoethanol, 2% SDS, 10% glycerol, 0.02% bromophenol blue) followed by heating for 5 min at 95°C. Protein lysates were subjected to SDS-PAGE and transferred onto PVDF membranes. Membranes were blocked in 5% nonfat powdered milk for 1 h at RT and incubated in primary antibodies overnight at 4°C. Membranes were washed 3 times for 10 min each in TBST(10 mM Tris/Cl, pH 7.6, 150 mM NaCl, 1% Tween 20), followed by incubation in goat anti-mouse or anti-rabbit secondary antibody coupled to horseradish peroxidase for 1 h at RT. Membranes were washed 3 times as above, followed by chemiluminescence detection with ECL reagents. Primary antibodies used were: anti-eIF3a (Novus, cat. NBP1-18891), anti-eIF3b (Bethyl, cat. A301-761A), anti-eIF3d (Bethyl, cat. A301-758A), anti-eIF3e (Bethyl, cat. A302-985A), anti-RPL7 (Abcam, cat. ab72550) and anti-RPS19 (Bethyl, cat. A304-002A).

### Sucrose density gradient centrifugation

Total lysate was prepared by scraping 6 × 10^7^ cells into 0.5 ml hypotonic buffer (5 mM Tris-HCl, pH 7.5, 2.5 mM MgCl_2_, 1.5 mM KCl and 1x protease inhibitor cocktail) supplemented with 100 μg/ml cycloheximide (CHX), 1 mM DTT, and 100 units of RNAse inhibitor. Triton X-100 and sodium deoxycholate were added to a final concentration of 0.5% each, and samples were vortexed for 5 seconds. Samples were centrifuged at 16,000 g for 7 min at 4°C. Supernatants (cytosolic cell extracts) were collected and absorbance at 260 nm was measured. Approximately 10-15 OD_260_s of lysate was layered over 10% – 50% cold sucrose gradients in buffer (200 mM HEPES-KOH, pH 7.4, 50 mM MgCl_2_, 1 mM KCl, 100 μg/mL CHX and 1x Pierce™ protease inhibitor). Gradients were centrifuged at 39,000 rpm in a Beckman SW28 rotor for 2 h at 4°C. After centrifugation, 14 equal-sized fractions (0.75 mL/fraction) were collected and analyzed through UV detection. For immunoblotting, fractions were mixed with SDS sample buffer.

### Pulsed SILAC sample preparation

MCF-10A cells were grown in DMEM: F12 (SILAC standard) containing light (^12^C, ^14^N) lysine and arginine supplemented with 5% dialyzed FBS and EGF, insulin and hydrocortisone as described above for 2 weeks. 1 × 10^7^ cells were treated with eIF3e or control siRNA for 48 h, followed by changing the light medium to heavy medium containing (^13^C, ^15^N) lysine and arginine. Cells were scraped in pre-cooled PBS and pelleted by spinning at 600 g for 5 min at 4°C. Cell pellets were resuspended in 0.5 mL 8 M urea lysis buffer (8 M urea in 0.2 M Tris/HCl, 4 mM CaCl_2,_ pH 8.0). The supernatant fractions were collected after centrifugation at 12,000 g for 15 min at 4 °C, and protein concentrations were determined by BCA assay (Pierce). Mass spectrometry for the pSILAC analysis was carried out by the National Center for Protein Science Beijing.

### Quantitative proteomics of polysomal fractions and purified 80S and eIF3 complexes

MCF-10A cell lysate was fractionated through a 35% sucrose cushion and the polysomal pellet was digested with RNAse I to dissociate polysomes into 80S monosomes. The 80S monosomes were further purified on a 10 – 50% sucrose density gradient. The total cell lysate, the digested polysomes and the 80S monosomes were analyzed by quantitative LC-MS/MS. For analysis of eIF3 complexes, 2 μg of anti-eIF3b antibody were pre-absorbed to 150 μl protein A + G magnetic beads for 2 h at 4 °C and incubated with gel filtration fractions enriched for eIF3 complexes overnight at 4 °C. Beads were washed with IP buffer (100 mM Tris, pH 7.5, 150 mM NaCl, 0.5% Triton X-100) 3 times for 5 min each Following trypsin digestion, peptide mixtures were re-dissolved in 0.1% formic acid in ultrapure water before being analyzed by HPLC coupled to a Q-Exactive mass spectrometer (Thermo Fisher Scientific) operated in positive data-dependent acquisition mode. Protein identification and quantitation were automatically performed by Thermo Proteome Discoverer (PD 1.4.0288) software against UniProt human protein database release 2016_09. Precursor ion mass tolerance was 10 ppm; fragment ion mass tolerance was 0.5 Da. The FDR of protein and peptide was 0.01. The normalized spectrum abundance factor (NSAF) was calculated by a custom Pearl script. The mass spectrometry data were submitted to PeptideAtlas (PASS01378).

### Transmission electron microscopy

Cell pellets after trypsinization corresponding to 1×10^7^ cells or fresh tissues removed from mice and dissected into pieces of a volume of ∼1 mm^3^ were washed with sodium cacodylate buffer pre-cooled to 4 °C (1 mM CaCl_2_ in 0.1 M sodium cacodylate buffer, pH 7.4) and immersed in pre-cooled fixation buffer (2.5% glutaraldehyde, 2.4% formaldehyde, and 1 mM CaCl_2_ in 0.1 M sodium cacodylate buffer, pH 7.4). Samples were fixed overnight at 4°C, washed with sodium cacodylate buffer three times and post-fixed in OsO4 (1% OsO4, 0.1% potassium ferrocyanide, 1 mM CaCl_2_ in 0.1 M sodium cacodylate buffer, pH 7.4) for 1.5 h at 4°C. Samples were dehydrated in 30% and 50% ethanol and stained in 70% uranyl acetate for 4 h, followed by serial dehydration in 70%, 90% and 100% ethanol. Samples were embedded in Spurr’s resin, sectioned into 75 nm slices, and stained with lead citrate for 10 min. Images were taken on a Tecnai Spirit BioTwin TEM.

To quantify mitochondrial area in muscle of wildtype and eIF3e^+/-^ mice, TEM micrographs of muscle in the longitudinal orientation at a magnification of ×13,000 were used. Mitochondrial size measurements were obtained using Image J (version 1.42q, National Institutes of Health, Bethesda, MD) by manually tracing only clearly discernible outlines of muscle mitochondria on TEM micrographs.

### Autophagic flux assay

1 × 10^6^ MCF-10A cells were simultaneously transfected with 2 μg pDest-mCherry-EGFP-LC3B and with eIF3e or control siRNA for 48 h using Lipofectamine 2000 transfection reagent. Prior to imaging, cells were treated with 0.2 µM bafilomycin A1 for 12 h, washed with Hank’s buffer, and live cells were imaged by confocal microscopy (Zeiss Exciter 5).

### Measurement of oxygen consumption rate

The Seahorse XFe96 metabolic flux analyzer was used for measuring oxygen consumption rate (OCR). Wells of the utility plate were filled with 200 μL of XF Calibrant and placed at 37 °C at atmospheric CO_2_ overnight before seeding 15,000 MCF cells in complete medium. After 12 – 24 hours, cells were washed with 100 μL XF Base medium (Seahorse Bioscience 102353-100) which has been adjusted to pH 7.4 at 37 °C with 0.1M NaOH. 175 μL XF Base medium was added to all 96 wells and the plate was placed at 37 °C in an incubator at atmospheric CO_2_ for 60 min. 1 μM oligomycin, 0.5 μM carbonyl cyanide 4-(trifluoromethoxy)phenylhydrazone (FCCP) and 1 μM of AA/rotenone were added to the injection wells and assays were run according to standard instrument protocol. The oxygen consumption rate was normalized to protein concentration.

### Generation of eIF3e knockout mice

eIF3e conditional knock-out mice crossed with FVB-Tg(Ddx4-cre)1Dcas/JNju mice which were purchased from Shanghai Biomodel Organism Science & Technology Development Co., Ltd. to breed eIF3e^+/-^ heterozygous knock-out mice in which exon2 of eIF3e in one of the DNA strand was floxed. And self-cross eIF3e^+/-^ heterozygous knock-out mice to obtain more eIF3e^+/-^ heterozygous knock-out mice. Genotyping was done by PCR using tail DNA as template. Mice tail were lysed in 500 μl lysis buffer (100 mM Tris, pH 8.0, 200 mM NaCl, 0.2% SDS, 5 mM EDTA, and 75 μg/μL PK enzyme) overnight (6 h) at 55°C, then centrifuged at 12,000 g for 10 min at RT to recover the supernatant. An equal volume of isopropanol was added followed by gentle shaking and centrifugation at 12,000 g for 10 min at RT. The DNA pellet was dissolved in 50 μl of ddH_2_O. eIF3e genotypes were determined by performing PCR using 100 ng purified genomic DNA as a template and the following primers: forward primer (5L-GGTGTGGAGAAGAAGAGAAGGT-3L) and reverse primer (5L-GCAGGGACAAAGAGTGGAACA-3L). All mice were in a C57BL/6J strain background.

### Grip strength assay

Forelimb grip strength of mice was measured with the Ugo Basile 47200 Grip-Strength Meter. Each mouse was tested five times and the peak force was recorded. Mice were measured at ages of 2, 3, or 4 months as indicated in Figure 6.

### NADH-tetrazolium reductase assay to measure complex I activity in skeletal muscle

The gastrocnemius and quadriceps muscles were isolated from mouse hind limbs and dehydrated in a 30% sucrose solution and then sectioned in a constant cold section box (Leica CM 3000, Germany) at −20 °C. 8-μm-thick frozen cross sections were fixed in pre-cooled acetone for 30 min at −20 °C. After washing 3 times for 3 min each in PBS, sections were stained by a nicotine amide adenine dinucleotide-tetrazolium reduction reaction containing 0.8 mg/ml NADH, 1 mg/ml nitrotetrazolium blue chloride and 10% DMSO in 0.2 M Tris, pH 7.5 for 6 minutes at 37°C. The reaction was terminated by washing with water. The images of muscle sections were observed by light microscopy (Olympus BX53, Japan).

### Uphill treadmill running test

Mice were acclimated on the treadmill (Ugo Basile 47300) tilted at a 10% slope at a speed of 20 m/min for 30 min on 3 constitutive days before the test day. After acclimation, mice were run at 10 m/min for 10 min, and speed was increased by 2 m/min every 1 min until mice were exhausted. Distance run until the mice failed to climb the treadmill within 5 s despite electric stimulation was recorded.

### Preparation of ribosome footprint libraries

1×10^7^ MCF-10A cells were lysed with polysome buffer for 10 min on ice, then 16,000 g spin for 10 min at 4° C to recover lysates (Ingolia et al., 2012). After clarification, 5 OD_260_ units of lysates were treated with 450 units of RNaseI (Ambion) for 45 min at RT. Ribosome protected fragments (RPFs) were isolated by chromatography on a Sephacryl S400 spin column (GE, MicroSpin S-400 HR, 27514001) according to manufacturer’s instructions. mRNAs and RPFs were isolated with TRIzol reagent. The rRNA removal for total mRNAs and RPFs was done using Ribo-Zero Gold rRNA removal kit (Illumina, cat, MRZG12324). RPFs were size-selected by 15% denaturing PAGE and cutting out the gel region corresponding to a size of 26 – 34 nucleotides. An oligonucleotide adaptor was ligated to the 3’ end of total mRNAs and RPFs, followed by reverse transcription, circularization and PCR amplification. cDNA libraries were quality tested by running on an Agilent 2100 Bioanalyzer and subjected to 50-bp single end sequencing on an Illumina Hiseq 2000 instrument.

### Selective ribosome profiling

2 μg of anti-eIF3b (Bethyl, cat. A301-761A) or anti-eIF3e (Bethyl, cat. A302-985A) antibody were pre-absorbed to 150 μl protein A + G Sepharose beads and beads were blocked in 0.5% BSA for 0.5 hours at 4 °C. Beads were incubated overnight with RPFs prepared as described above (the RPF material recovered upon Sephacryl S400 chromatography) at 4 °C. Beads were washed with IP buffer (100 mM Tris, pH 7.5, 150 mM NaCl, 0.5% Triton X-100) 3 times for 5 min each), and associated RNA was isolated with TRIzol reagent. The cDNA library was constructed and sequenced as described above. All RNAseq data were submitted to Gene Expression Omnibus (GSE131074).

### Pre-processing of the ribosome profiling and mRNA-seq data

The human reference genome assembly (GRCh38) and the annotation file downloaded from the Ensembl genome browser were used for all analyses. The pre-processing procedure of the ribosome profiling and mRNA-seq data has been described previously (Xiao et al., 2016). FastQC (http://www.bioinformatics.babraham.ac.uk/projects/fastqc/) was used for quality control and the adaptor sequence (CTGTAGGCACCATCAAT) in the raw reads of both ribosome profiling and mRNA-seq was trimmed using the Cutadapt program. Reads with length between 25-35 nt were used for ribosome profiling analyses. Low-quality reads with Phred quality scores lower than 25 (>75% of bases) were removed using the fastx quality filter (http://hannonlab.cshl.edu/fastx_toolkit/). Then sequence reads originating from rRNAs were identified and discarded by aligning the reads to human rRNA sequences downloaded from GenBank (https://www.ncbi.nlm.nih.gov/genbank/) using Bowtie 1.1.2 with no mismatch allowed. The remaining reads were mapped to the genome using STAR (Dobin et al., 2013) with the following parameters: --runThreadN 8 --outFilterType Normal --outWigType wiggle -- outWigStrand Stranded --outWigNorm RPM --alignEndsType EndToEnd -- outFilterMismatchNmax 1 --outFilterMultimapNmax 1 --outSAMtype BAM SortedByCoordinate --quantMode TranscriptomeSAM GeneCounts --outSAMattributes All.

The RNA-seq reads for each gene were counted with HTSeq-count in intersection-strict mode and the read counts of RPF were calculated with a custom script written by Xiao (Xiao et al., 2016). The translation efficiency (TE) of each gene was calculated by dividing the read count of mRNA by that of RPF. All other analyses were performed using custom scripts written with Python 3.6 and R 3.4.3.

### Assessment of 3-nt periodicity and offset determination

3-nt periodicity of the RPF reads aligned by their P-sites was used as a strong evidence of activate translation (Ingolia et al., 2009). We used the RiboCode program (Xiao et al., 2018) to assess the 3-nt periodicity of all Ribo-seq samples.

### Meta-analysis of ribosome profiling data

The read counts at each codon of the transcripts were calculated using a custom Python script. As for each transcript, the read counts vector is written as:

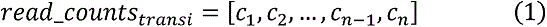

Where ci is the read count at codon position i in the transcript, and n represents the length of the ORF in unit of codon. The read counts at each position were normalized by the mean value of each read counts vector without the first 30 codons to exclude the accumulated reads around the start codon. For genes with multiple transcript isoforms, only the longest isoform was used when parsing the genome annotation files.

For the metagene analyses at the global scale, transcripts with more than 300 codons were used. Low level transcripts (raw counts fewer than 128 or normalized counts less than 64) were discarded. However, when specifically studying the metagene spectrum between the codons 25 and 75, the transcripts with ORF length longer than 150 codons and RPKM (reads per kilobase per million mapped reads) values larger than 10 were used. After the filtering steps above, all transcripts were lined up and the mean read density at each position was be calculated as:

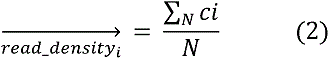

Where N is the numbers of transcripts retained, and ci represents the read density at codon i position. Finally, all mean read density at each position formed a vector that was then used for metagene plots with a moving-average method (size of window: 7 codons; step size: 1 codon).

To identify the mRNAs that contributed to the accumulated RPF reads between codons 25 - 75 upon eIF3E knockdown, we implemented a bootstrap-based procedure, which partitioned the range of codons 25 - 75 into 9 windows with 10 codons each (i.e., bin width = 10 codons, step width = 5 codons). Next, the transcripts that have higher (≥ 2-fold) read density in at least one of the 9 windows and are not lower than control in any other window when comparing the si-eIF3E cells to control knockdown cells were extracted and used for further analysis. Finally, 2683 transcripts were identified with ≥ 2-fold accumulation of ribosomes between codons 25 - 75.

### Calculation of ribosome polarity scores

To quantify global differences in the position of ribosomes along transcripts, we computed a polarity score for every gene as described previously (Schuller et al., 2017). The polarity at position i in a gene of length l is defined as follows:

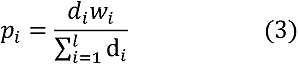

Where

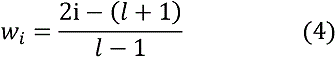

The *d_i_* and *w_i_* represent the ribosome density and normalized distance from the center of a gene at position i respectively. Polarity score for a gene is the total sum of pi at each position. Also, we excluded the first and the last 15 nt of coding sequences from our analyses. mRNAs with more than 64 reads in coding sequences were used for generating the polarity plots.

### Data analysis for selective ribosome profiling

Read counts along each transcript were computed as described in the section above (Meta-analysis of ribosome profiling data). Moving-average method was used to obtain read density at each position along a transcript (size of the window: 7 codons; step size: 1 codon). Enrichment ratios were calculated at each position (i.e., RPFs obtained from eIF3b- or eIF3e-associated ribosomes over RPFs obtained for all ribosomes). Remaining zeros at some positions were replaced with the number 1. Transcripts with a global difference in the positioning of eIF3b- or eIF3e-associated ribosomes relative to all ribosomes were identified by comparing their polarity scores (see section above). Specifically, an mRNA with a smaller polarity score based on the eIF3-bound RPF density than the score based on RPF density obtained by conventional ribosome profiling (polarity score difference <0) was defined as an “eIF3-80S 5’-enriched mRNA”. In total, 2543 such mRNAs were used for meta-analysis for eIF3b selective ribosome profiling. The 5204 mRNAs with a polarity score difference >0 were defined as “eIF3-80S 5’-depleted mRNAs”. For eIF3e selective ribosome profiling, 4623 mRNAs had a polarity difference <0, while 4179 mRNAs had a polarity difference >0. Only transcripts longer than 150 codons with RPKM in the coding region larger than 10 were used. Meta-analysis for the enrichment ratios was done as described above for conventional ribosome profiling.

### Hydrophobicity index

Hydrophobicity index (also called hydropathy index) of an amino acid is a number representing the hydrophobic or hydrophilic properties of its sidechain. The hydrophobicity index was download from AAindex (https://www.genome.jp/aaindex/) and the mean values at each position along the amino acid sequences were fed into a vector used for metagene analysis.

### Charges of amino acid

Amino acids with positive charges (Lys and Arg) were assigned a score of 1, amino acids with negative charge were assigned a score of −1, and neutral amino acids were assigned a score of 0. Proteins were aligned at the starting methionine and charge scores at each residue were computed and averaged across all proteins in the analysis. Average scores of all proteins were plotted.

### The local tAI

The local tAI (tRNA adaptation index) was computed as previously described (Tuller et al., 2010). That is, let *n_i_* be the number of tRNA isoacceptors recognizing codon i. Let tGCNij be the copy number of the jth tRNA that recognizes the ith codon and let Sij be the selective constraint on the efficiency of the codon-anticodon coupling. The absolute adaptiveness, Wi, for each codon i is

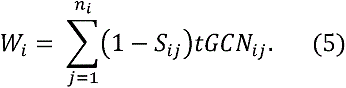

From the absolute adaptiveness we could get the relative adaptiveness value of codon i, namely wi, by normalizing the Wi values (dividing them by the maximum of all Wi values).

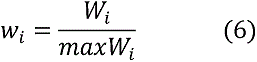

The classic tAI of a gene later would be calculated as the geometric mean of wi. In the present study, however, we intended to observe the tAI values at each position along transcripts.

Therefore, wi was treated as the local tRNA adaptiveness index for codon i, after which all local tAI values of each transcript were lined up and mean values at each position were computed and plotted. The copy numbers of each tRNA were downloaded from GtRNAdb (http://gtrnadb.ucsc.edu/) and the scores Sij for wobble nucleoside-nucleoside pairing were taken from Tuller et al. (Tuller et al., 2010).

### Kozak consensus context

Sequences containing from 10 nucleotide upstream to 8 nucleotide downstream of the start codon were extracted from transcript sequences and motif logos were generated by Weblogo3 (http://weblogo.threeplusone.com/create.cgi).

### uORF analysis

Translated uORF were identified and annotated using RiboCode as described (Xiao et al., 2018). A modified Wilcoxon signed rank test was used to assess the 3-nt periodicity of RPF reads mapped on different reading frames (f0, f1, f2) for a given candidate uORF. If the test resulted in a statistically significant 3-nt periodicity (p-value smaller than the cutoff, e.g. 0.05), the uORF was considered actively translated.

### Quantification of tRNA expression

The mature tRNA sequences (hg38) were downloaded from gtRNAdb (http://gtrnadb.ucsc.edu/) and appended with a CCA tail using a custom python script. The CAA tail was used as reference for mapping. Given sequence similarities between tRNA iso-acceptors, we used USEARCH (https://www.drive5.com/usearch/) for tRNA clustering to avoid possible multiple mapping. Bowtie (Langmead et al., 2009) was then used for tRNA mapping based on the total RNA-seq datasets. A GTF file was generated by a custom python script, and HTSeq (Anders et al., 2015) was used for tRNA quantification. DESeq2 (Love et al., 2014) was used for data normalization.

### Statistical analysis

All p values for the ribosome profiling data were calculated using t-tests based on custom R or Python scripts. Statistical analysis of the remaining datasets was performed with Microsoft Excel. Typically, data were averaged, standard deviations calculated, and statistical significance was assessed using the T.Test function assuming two-tailed distribution and unequal variance.

### Functional pathway analysis

For each given gene list, pathway and process enrichment analysis has been carried out with Metascape (metascape.org (Zhou et al., 2019)) with the following ontology sources: GO Biological Processes, GO Cellular Components, Canonical Pathways and CORUM. All genes in the genome were used as the enrichment background. Terms with a p-value < 0.01, a minimum count of 3, and an enrichment factor > 1.5 (the enrichment factor is the ratio between the observed counts and the counts expected by chance) were collected and grouped into clusters based on their membership similarities. According to the description at metascape.org, p-values were calculated based on the accumulative hypergeometric distribution, and q-values were calculated using the Benjamini-Hochberg procedure to account for multiple testing. To capture relationships between enriched functional terms, a subset was selected and rendered as a network plot, where terms with a similarity > 0.3 were connected by edges. Terms with the best p-values from each of 20 clusters were selected, and the network was downloaded as a Cytoscape (Cline et al., 2007) file. Each node represents an enriched term colored by its p-value.

---

**Supplementary Fig. 1.**
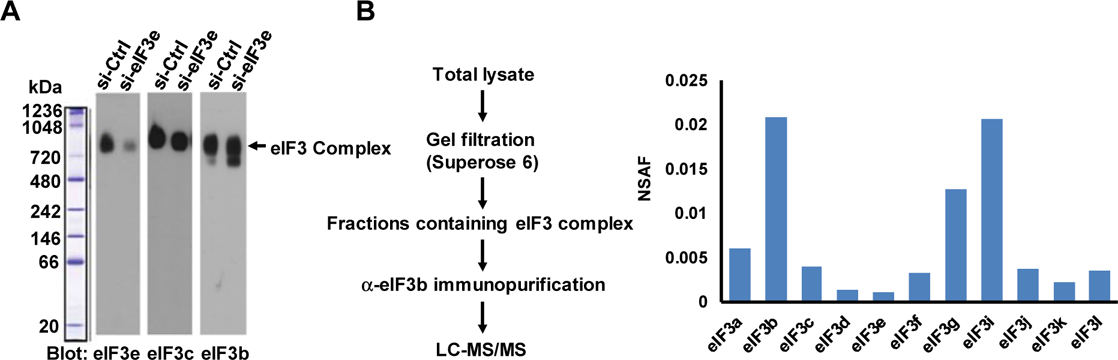
Effect of eIF3e knockdown on eIF3 holo-complex (related to Fig. 1) **A.** MCF7 cells were transfected with si-eIF3e or si-Control for 72 h, followed by separation of cell lysate by native PAGE and detection of eIF3 by immunoblotting with the indicated antibodies. eIF3e knockdown depletes eIF3e from the ∼800 kDa eIF3 complex, but other subunits, including eIF3b and eIF3c as well as the overall size of the complex are maintained. **B.** Demonstration that the ∼800 kDa complex shown in a) corresponds to the eIF3 holo-complex. MCF7 lysate was separated by gel filtration on a S300 column, and the fractions containing the ∼800 kDa complex were pooled and used for immunoprecipitation with eIF3b antibodies or normal rabbit IgG as control. The immunopurified material was analyzed by LC-MS/MS, and eIF3 subunits were quantified using normalized spectral abundance factors (NSAF). The data shows that the ∼800 kDa complex contains most eIF3 subunits, although subunits eIF3h and eIF3m were not identified. The IgG negative control did not pull down any eIF3 subunits.

**Supplementary Fig. 2.**
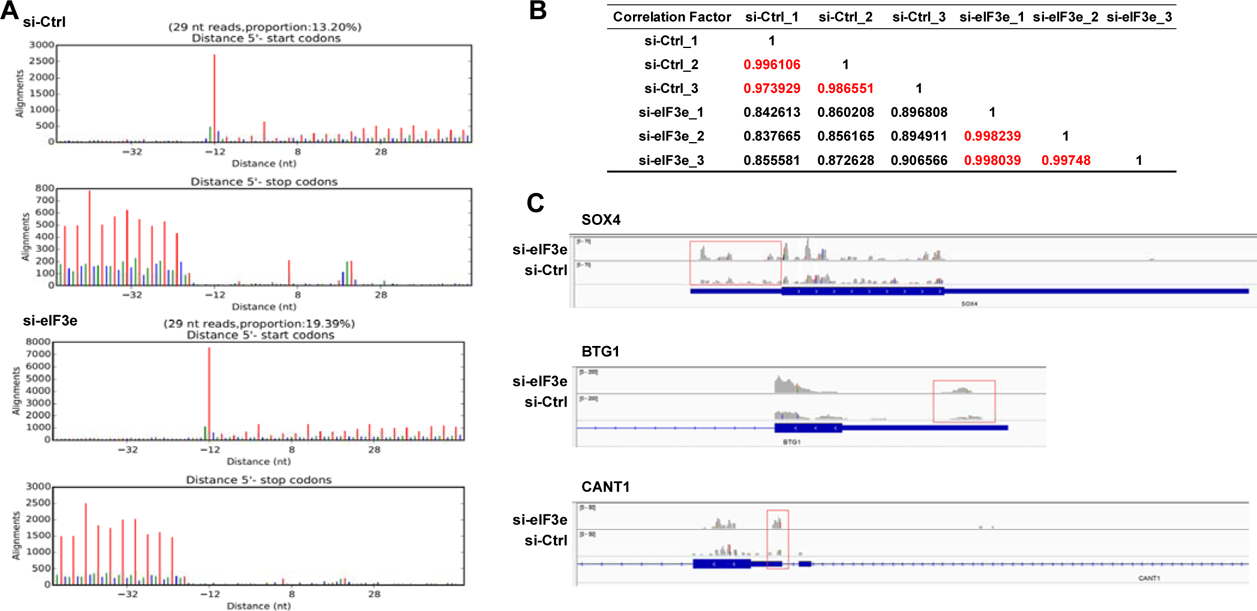
Quality and reproducibility of the ribosome profiling datasets (related to Fig. 1) **A.** RPF reads derived from si-Control and si-eIF3e samples and mapped to the human transcriptome reveal the 3-nucleotide periodicity typical of ribosome protected fragments. **B.** Correlation coefficients for the triplicate datasets obtained from si-Control and si-eIF3e samples. **C.** Ribosome densities in uORFs and cORFs of the indicated mRNAs in si-Control and si-eIF3e samples.

**Supplementary Fig. 3.**
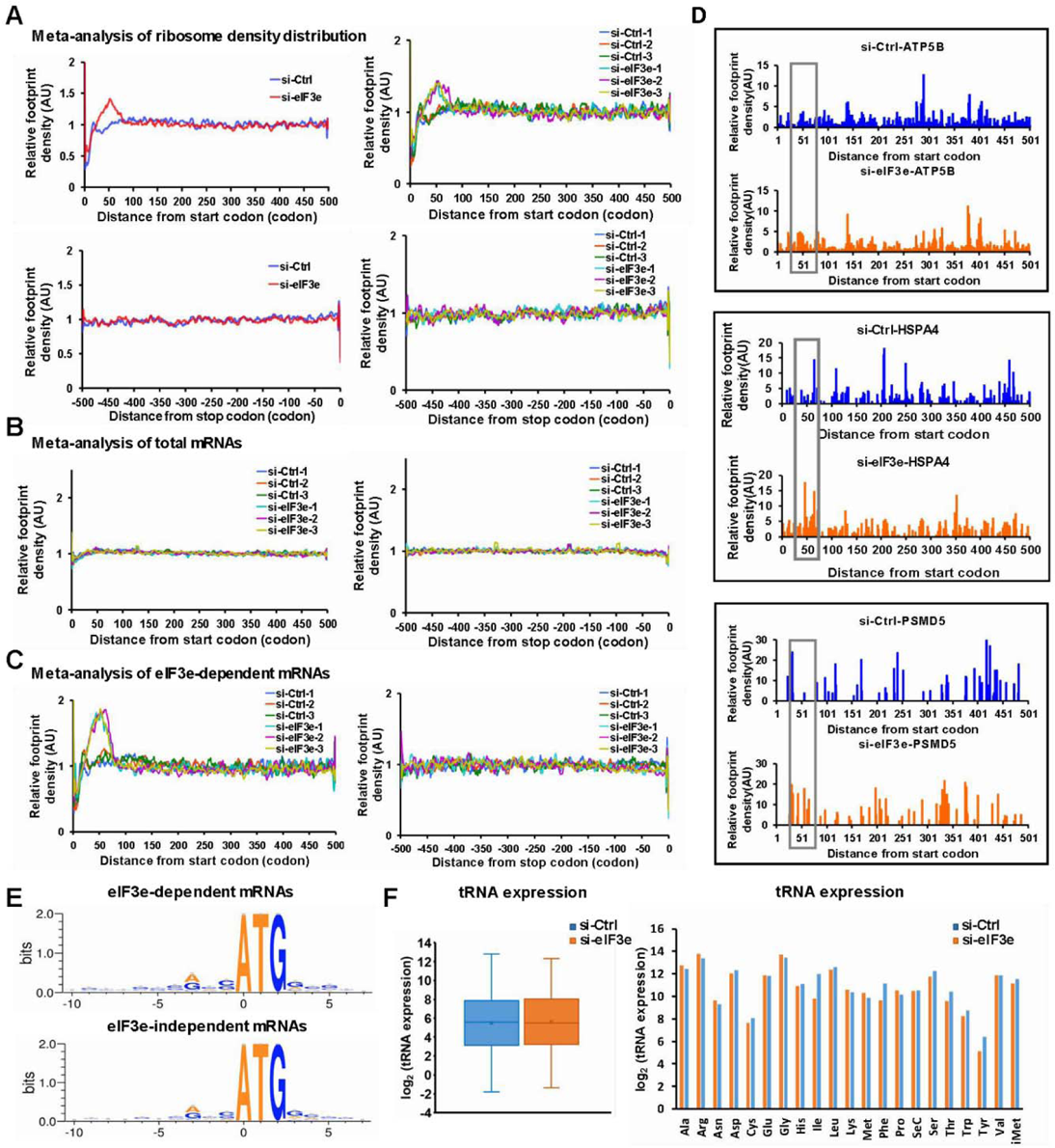
Evidence for ribosomal pausing between codons 25 and 75 (related to Fig. 2) **A.** Meta-analysis of ribosome density distribution for the total datasets after filtering (see Methods for details). The left panel shows averages from triplicate si-Control and si-eIF3e datasets, the right panel shows the three individual datasets. Top panels show the region from the start codon to codon 500, the bottom panel shows the last 500 codons before the stop codon. **B.** Same meta-analysis as in a) performed on the total mRNA datasets, which do not show a peak between codons 25 - 75. **C.** Meta-analysis of ribosome density distribution for the set of 2683 mRNAs which depend on eIF3e for efficient early translation elongation. Individual replicates are shown for the regions 500 codons downstream of the start codon and upstream of the stop codon. **D.** Examples of individual mRNAs displaying selective ribosome accumulation in the region between 25-75 codons. **E.** Comparison of the Kozak consensus context between 2863 eIF3e-dependent and 5320 eIF3e-independent mRNAs. **F.** tRNA levels in MCF-10A cells after knockdown of eIF3e relative to si-Control.

**Supplementary Fig. 4.**
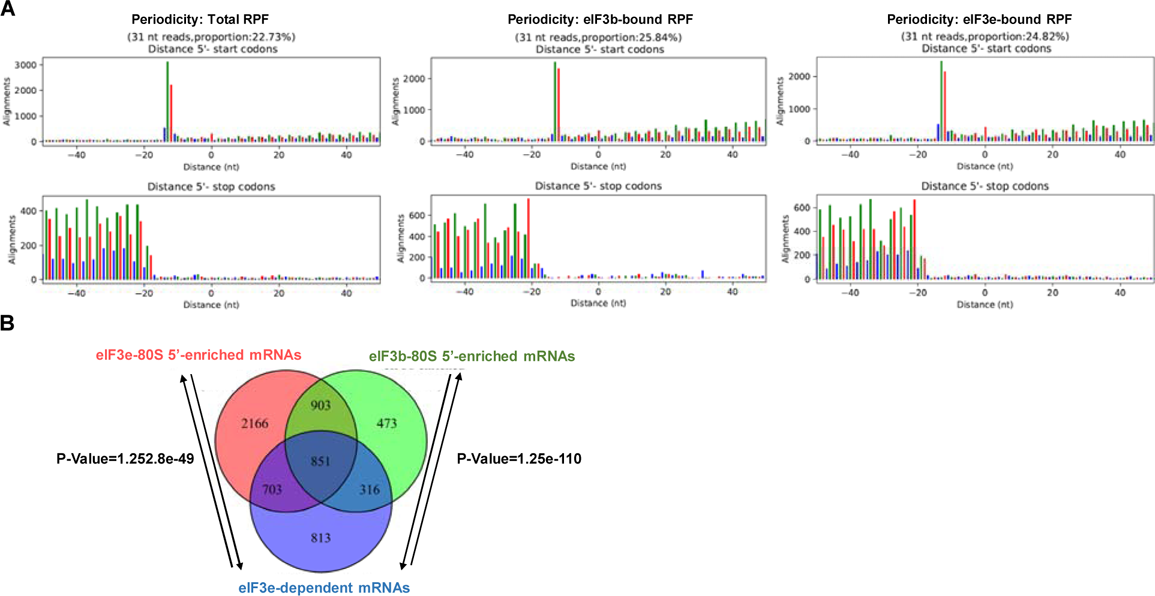
Characteristics of the eF3b and eIF3e selective ribosome profiling datasets (related to Fig. 5) **A.** 3-nucleotide periodicity of the RPFs derived from all ribosomes and from eIF3b- and eIF3e-associated ribosomes. **B.** Overlap in the set of 2683 eIF3-dependent mRNAs and the sets of 2543 (eIF3b) and 4623 (eIF3e) mRNAs with increased density of eIF3-associated ribosomes (“eIF3-80S 5’-enriched mRNAs”).

**Supplementary Fig. 5.**
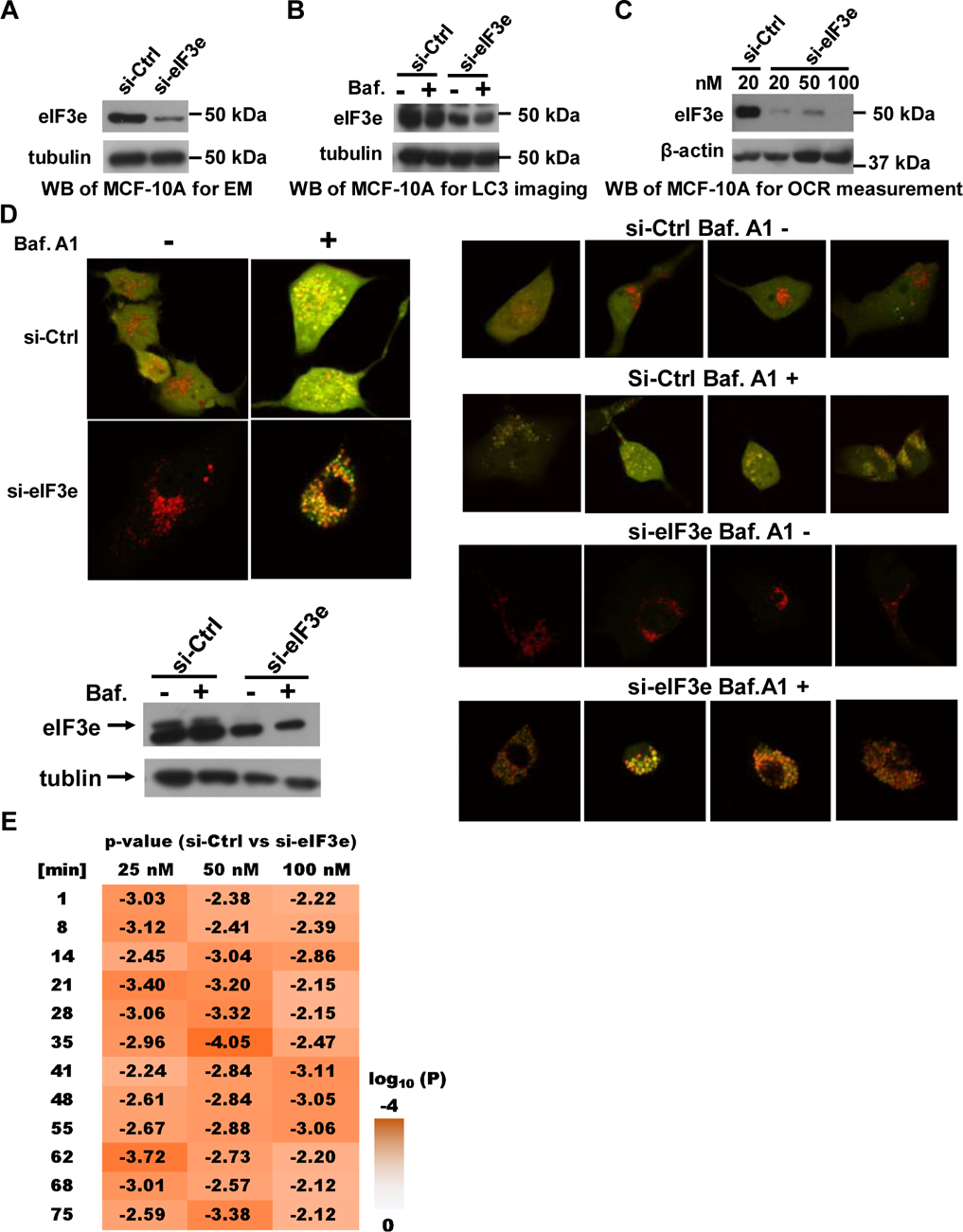
Confirmation of eIF3e knockdown efficiencies for experiments shown in Fig. 6. **A.** Immunoblot to confirm the knockdown of eIF3e in MCF-10A cells used for electron microscopy (Fig. 6A). **B.** Immunoblot to confirm the knockdown of eIF3e in MCF-10A cells used for live cell imaging of ectopically expressed mCherry-EGFP-LC3B fusion protein (Fig. 6B). **C.** Immunoblot to confirm the knockdown of eIF3e in MCF-10A cells used for measurement of oxygen consumption rate (Fig. 6C). **D.** Independent repeat of the experiment in Fig. 6B, showing additional examples of si-Control and si-eIF3e cells expressing mCherry-EGFP-LC3B with or without bafilomycin A1 (Baf. A1) treatment. The immunoblot confirms the knockdown of eIF3e in MCF-10A cells used in this experiment.

**Supplementary Fig. 6.**
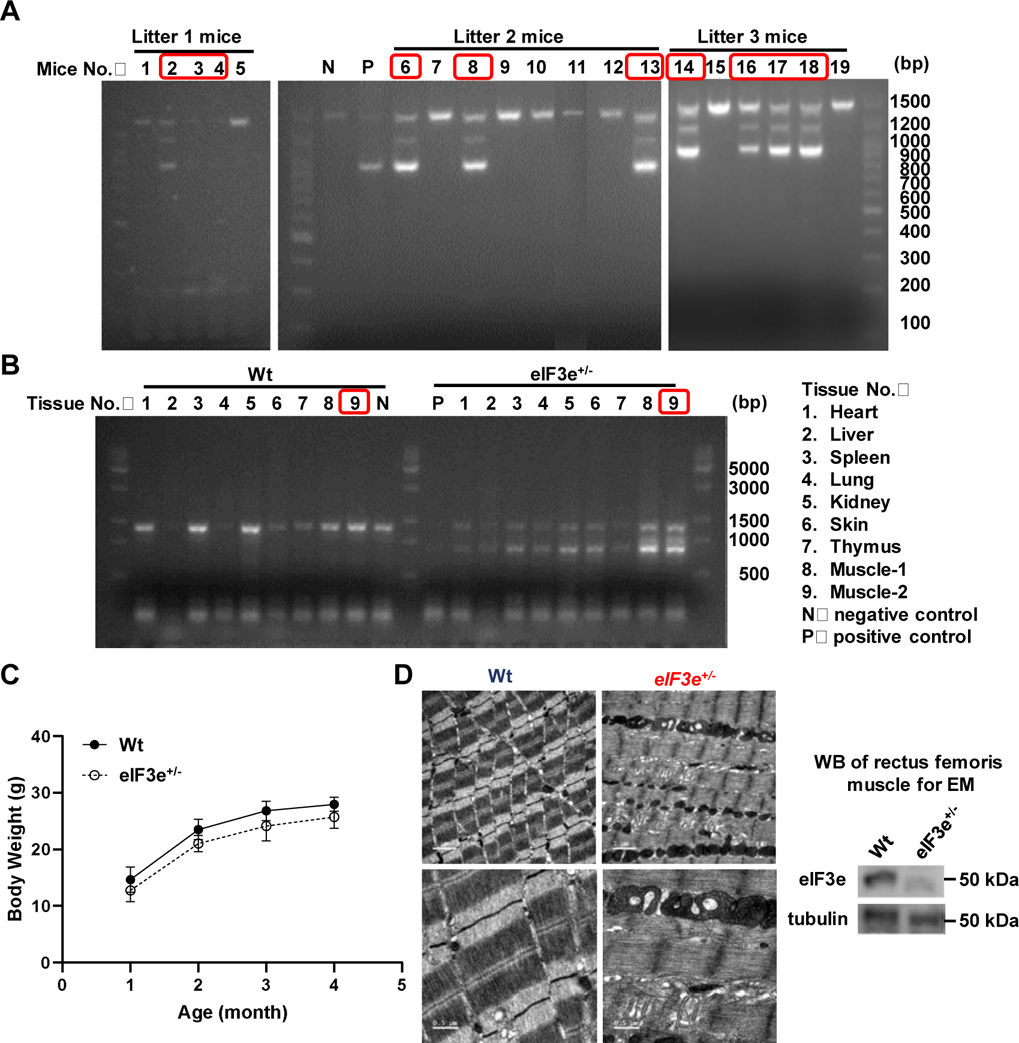
Genotyping of eIF3e+/-mice and muscle phenotype (related to Fig. 6) **A.** PCR genotyping using tail DNA from three different litters of mice from crosses of eIF3^+/-^ females and males. Heterozygous eIF3^+/-^ mice resulting from the crosses and used in this study are highlighted. **B.** PCR genotyping of DNA from the indicated tissues confirms heterozygous knockout of eIF3e. The muscle tissue used for electron microscopy are highlighted. **C.** Body weights of eIF3e^+/-^ mice. **D.** Electron micrographs of rectus femoris muscle from wildtype and eIF3^+/-^ heterozygous knockout mice. The immunoblot confirms the decrease of eIF3e in the muscle tissue shown.

**Supplementary Fig. 7.**
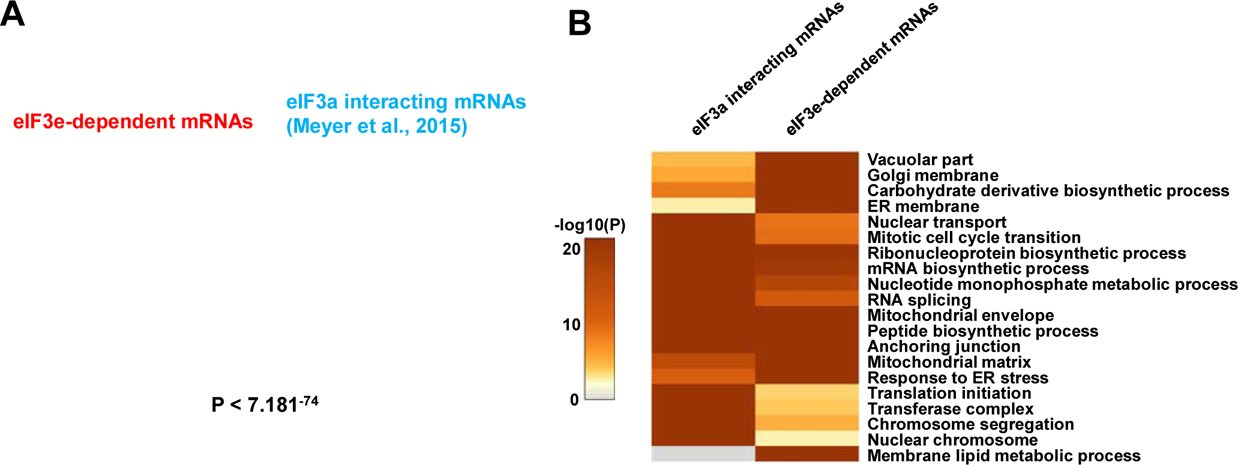
Overlap between eIF3-dependent and interacting mRNAs (related to Fig. 7) **A.** Venn diagram showing the intersection between the list of 2683 mRNAs identified by ribosome profiling to require eIF3e for early translation elongation (“eIF3e-dependent mRNAs”) and the list of mRNAs found to directly interact with eIF3a (Meyer et al., 2015). **B.** Overlap in the enrichment of functional terms in the set of 2683 eIF3e-dependent mRNAs and the set of mRNAs found to directly interact with eIF3a (Meyer et al., 2015).

